# Circulating osteopontin released by injured kidneys causes pulmonary inflammation and edema

**DOI:** 10.1101/2021.07.20.452998

**Authors:** Fatima Zohra Khamissi, Liang Ning, Eirini Kefaloyianni, Hao Dun, Akshayakeerthi Arthanarisami, Amy Keller, Jeffrey J. Atkinson, Wenjun Li, Brian Wong, Sabine Dietmann, Kory Lavine, Daniel Kreisel, Andreas Herrlich

## Abstract

Multiorgan failure is devastating, and its mechanisms and mediators are not clear. Tissue injury in one organ appears to trigger disease in remote organs. Kidney and lung are frequently affected, such as when acute kidney injury (AKI) causes acute lung injury (ALI), a frequent clinical condition with high mortality. Here we identify factors secreted from the injured kidney that cause acute lung injury. We developed a murine model mimicking the generation of respiratory failure following acute kidney injury. To identify interorgan crosstalk mediators involved, we performed scRNAseq of mouse kidneys and lungs after AKI. We then applied ligand-receptor (L-R) pairing analysis across cells residing in kidney (ligands) or lung (receptors) to identify kidney-released circulating osteopontin (OPN) as a novel mediator of AKI-induced ALI (AKI-ALI). OPN release very early after AKI largely from tubule cells triggered neutrophil and macrophage infiltration into lungs associated with endothelial leakage, interstitial edema, and functional impairment. Pharmacological or genetic inhibition of OPN prevented AKI-ALI. Transplantation of ischemic *wt* kidneys into *wt* mice caused AKI-ALI, while transplantation of ischemic OPN-global-knockout kidneys failed to induce lung endothelial leakage and AKI-ALI, identifying circulating kidney-released OPN as sufficient to cause AKI-ALI *in vivo*. We show that AKI in humans results in elevations in OPN levels in the serum. Increased serum OPN levels in patients with multiorgan failure have been shown to positively correlate with reduced kidney function, respiratory failure, and mortality. Thus, our results identifying OPN as a mediator of AKI-ALI may have important therapeutic implications in human AKI-ALI and multiorgan failure.

## Introduction

Homeostasis in multicellular organisms and the response to its disturbance by injury is coordinated by cell-cell communication aimed at maintaining and reestablishing homeostasis^1–3^. Studies on cellular functions in health and disease to date mostly focus on understanding cell-cell communication and its mediators within a given organ or diseased tissue^4^. However, mediators of interorgan crosstalk between cells in different organs or tissues that drive multiorgan failure in complex diseases with high mortality, their cellular sources and target cells are largely unknown. Clinically important examples of such interorgan crosstalk include acute kidney injury (AKI)-induced acute lung injury (ALI) (AKI-ALI) with respiratory failure^5–7^, acute lung injury and its most severe form acute respiratory distress syndrome (ARDS) followed by AKI^8–10^, or lung transplant-induced AKI^11^.

AKI is a common problem in the human population and develops in 2-5% of patients during hospitalization, in 50% of intensive care unit (ICU) patients, and in about 20% of kidney transplant patients within the first 6 months after transplantation^11,12^. Irrespective of its cause, AKI alone has a 15-30% mortality, which rises to 60-80% when AKI induces remote secondary organ complications (multiorgan failure), in particular AKI-ALI^5,7,12^. Molecular mechanisms and mediators of AKI-ALI are not yet well understood (reviewed in^5,7,13,14^). Candidate AKI-ALI mediators were largely derived from global gene knockout studies or from the use of systemic interventions. Such studies implicated interleukin-6 (IL6) and tumor-necrosis-factor (TNF), as well as IL10 released from splenic CD4 T-cells^15–18^ and the alarmin high mobility group box-1 (HMGB1)^15^ as putative mediators. For IL6 or TNF, as examples, it is known that AKI upregulates their expression in the kidney, and elevated serum levels of IL6 and TNF protein after AKI were generally interpreted to result from release of these mediators by the injured kidney. However, kidney cell-type specific knockouts of reported AKI-ALI mediators have not been tested in AKI-ALI models. Thus, it has not been conclusively shown that only or mainly the kidney serves as a source of these mediators after kidney injury. Further, which lung cell types or immune cells are targeted by these mediators, is largely unknown, except for a possible role of the TNF receptor TNFR1 in apoptosis of lung endothelial cells during AKI-ALI^16,17^.

Hormones, growth factors, chemokines, cytokines, neurotransmitters, and other secreted proteins (in the following referred to as ligands) act in cell-cell communication in an autocrine/paracrine fashion locally, but may have distant endocrine effects if released into the circulation. These ligands could be relevant for interorgan crosstalk, such as in the case of IL6 and TNF. We therefore chose to concentrate first on soluble ligands expressed in the kidney with the idea that these and the responding receptors expressed in lung cells will lead us to identify relevant mediators of interorgan communication. Ligand-induced cognate receptor activation in receiving cells generally results in altered gene expression and phenotypic changes. To date, Ligand–Receptor (LR) pairing analysis using bulk or single-cell/single nucleus mRNAseq (scRNAseq or snRNAseq) gene expression datasets has been used to infer cell-cell communication within a local tissue or organ^1,4^. Here, we present to our knowledge the first ligand-receptor (LR) pairing analysis across different organs to successfully determine a mediator of interorgan crosstalk in multiorgan failure *in vivo*. Our studies identify kidney tubule cell-released circulating osteopontin (OPN) as a key mediator of AKI-induced remote acute lung injury with respiratory failure (AKI-ALI).

## Results

### Ischemic acute kidney injury (AKI) causes severe acute lung injury (ALI)

Aiming at identifying communication between an injured tissue and other organs, we established a model of multiorgan failure. We chose to injure the kidney and study remote effects on the lung. We subjected C57BL/6 mice to bilateral renal ischemia-reperfusion-injury (IRI) and analyzed remote acute lung injury (AKI-ALI) (**Experimental Scheme Figure 1A**). Kidney injury caused highly elevated blood-urea-nitrogen (BUN) levels on Day 1 after AKI, indicating severe kidney failure, with slow improvement to about 50% over Days 3-5 (**Figure 1B**). Consistent with this, serum creatinine was significantly elevated on Day 1 and remained elevated at lower levels over Days 3-5 (**Figure 1C**). Severe injury to lungs and inflammation (AKI-ALI) followed with the same kinetics, as determined by expansion of interstitial spaces, increased cellularity, and interstitial edema (**Figure 1D**). Alveolar wall thickness increased over sham levels by 75% on Day 1, signaling impediment of oxygenation (**Figure 1E**). Neutrophils and interstitial macrophages accumulated in the lung, peaked on Day 1 after AKI, and were still significantly elevated by Day 5. Interstitial macrophages (IM) that accumulated in lungs after AKI are small and CD68low. Alveolar macrophages (CD68high) did not change in numbers (**Figure 1F**). We confirmed these results by mass cytometry (CyTOF) of lung single-cell preparations from sham and AKI mice (**Figure 1G**). The functional consequence of these changes was a significant impairment in oxygen exchange on Day 1 after AKI (**Figure 1H**). Our mouse data closely resemble what can be observed in patients with multiorgan failure after AKI^5,7,10^.

**Figure 1:**
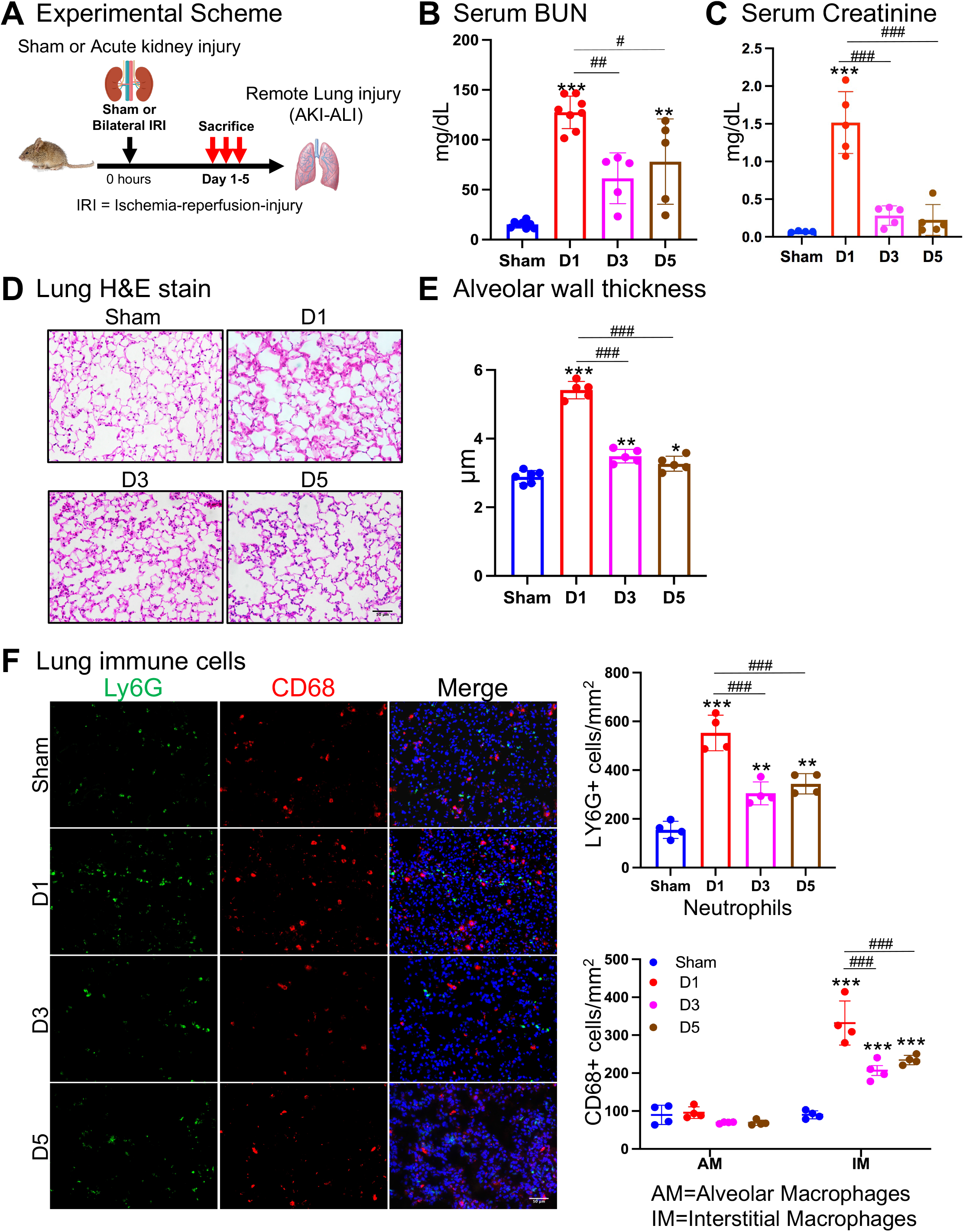
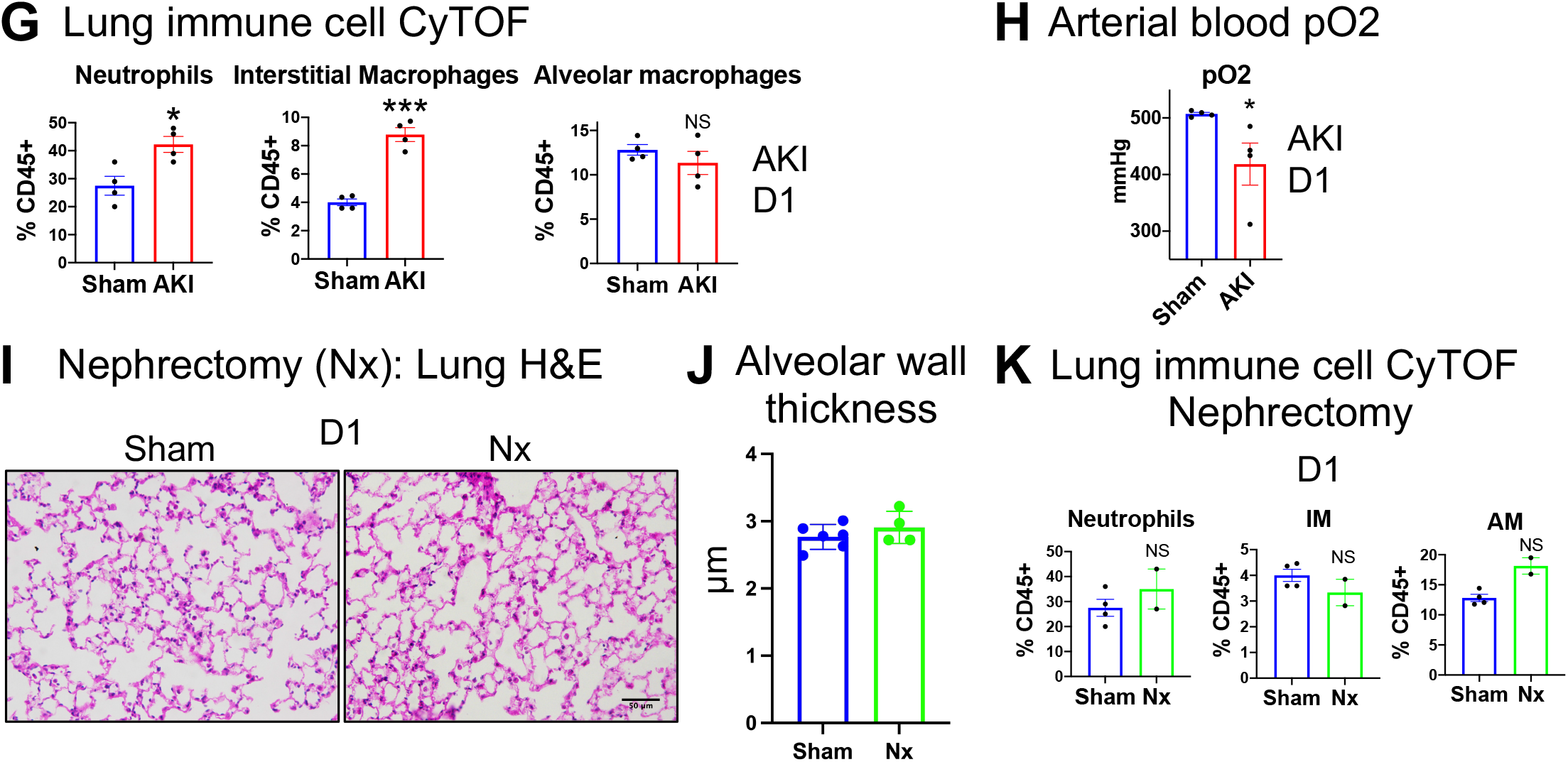
Ischemic acute kidney injury (AKI) causes severe acute lung injury (ALI) (**A.**) Experimental scheme: AKI → AKI-ALI model sham, Day 1, 3, 5 after AKI, (**B.**) Serum BUN values after Sham or AKI Day 1-5, (**C.**) Serum creatinine values after Sham or AKI Day 1-5, (**D.**) H&E stain lung after Sham or AKI Day 1-5, (**E.**) Alveolar thickness measurements after Sham or AKI Day 1-5, (**F.**) Lung neutrophils (Ly6G^+^, green), alveolar (CD68^high^, large, red) and interstitial macrophages (CD68^low^, small, red) and quantification after Sham or AKI Day 1-5. DAPI stain (blue) was used to visualize nulei, (**G.**) CyTOF: Lung neutrophils (CD45^+^, Ly6G^+^), alveolar macrophages (CD45^+^, CD68^high^, Siglec-F^-^) and lung interstitial macrophages (CD45^+^, CD68^low^, Siglec-F^-^) Day 1, (**H.**) Arterial blood oxygen partial pressure after Sham or AKI Day 1, (**I.**) Lung H&E after Sham or nephrectomy (Nx) Day 1, (**J.**) Alveolar thickness after Sham or nephrectomy (Nx) Day 1, (**K.**) CyTOF: Lung neutrophils (CD45^+^, Ly6G^+^), alveolar (CD45^+^, CD68^high^, Siglec-F^-^) and lung interstitial macrophages (CD45^+^, CD68^low^, Siglec-F^-^) were quantified after Sham or nephrectomy (Nx) Day 1. n=4-8 animals per measurement, *** p<0.001, ** p<0.01, * p<0.05, ### p<0.001, ## p<0.01, # p<0.05.

To exclude that the observed AKI-ALI phenotype is simply the result of accumulation of toxic waste products or failure to excrete an existing potentially detrimental molecule because of lack of glomerular filtration, we examined mice for AKI-ALI on Day 1 after simple bilateral nephrectomy, without reperfusion injury. Sham-operated mice or animals that underwent bilateral nephrectomies showed no differences in terms of lung interstitial spaces and cellularity (**Figure 1I**), alveolar wall thickness (**Figure 1J**) or lung immune cell numbers as assessed by CYTOF on Day 1 after nephrectomy (**Figure K**). These findings indicate that AKI-ALI does not develop simply due to a failure to excrete a potentially detrimental molecule.

### Single cell RNAseq of kidney and lung in setting of AKI-induced ALI (AKI-ALI)

As an entrance into identifying relevant cellular changes after AKI, we analyzed cellular gene expression profiles by single-cell mRNA sequencing (scRNAseq) of kidney and lungs isolated on Day 1 after sham-operation or bilateral renal ischemia-reperfusion-injury AKI (**Experimental Scheme Figure 2A**). Serum BUN levels were significantly elevated in injured mice on Day 1 after AKI (**Figure 2B**), indicating severe AKI. Analysis of scRNAseq gene expression profiles revealed the presence of all known non-immune and immune cell-types of the kidney and lung. Shown are Seurat objects that combine the data for sham and injury, as well as dotplots identifying standard marker genes used to identify cell types^18,19^, for either kidney (**Figure 2C+D**) or lung (**Figure 2E+F**). To monitor specific changes between sham and injury, we replotted our scRNAseq data separately for sham and injury of kidney or lung. Based on number of cells captured in each population (Seurat cluster), the injured kidney in comparison to sham showed easily detectable and expected dynamics in the immune cell compartment, with strong enrichment of neutrophils and monocytes in the injured sample (**Figure 2G**). Similar, but somewhat more subtle changes were detected in the same immune cells in the remotely injured lung. We detected a strong increase in the number of neutrophils and a small increase in monocytes in remotely injured lung tissue (**Figure 2H**). Changes in lung interstitial macrophage numbers between sham and injury were less appreciable at this level of our scRNAseq analysis, although we detected them in large numbers (**Figure 2G+H**).

**Figure 2:**
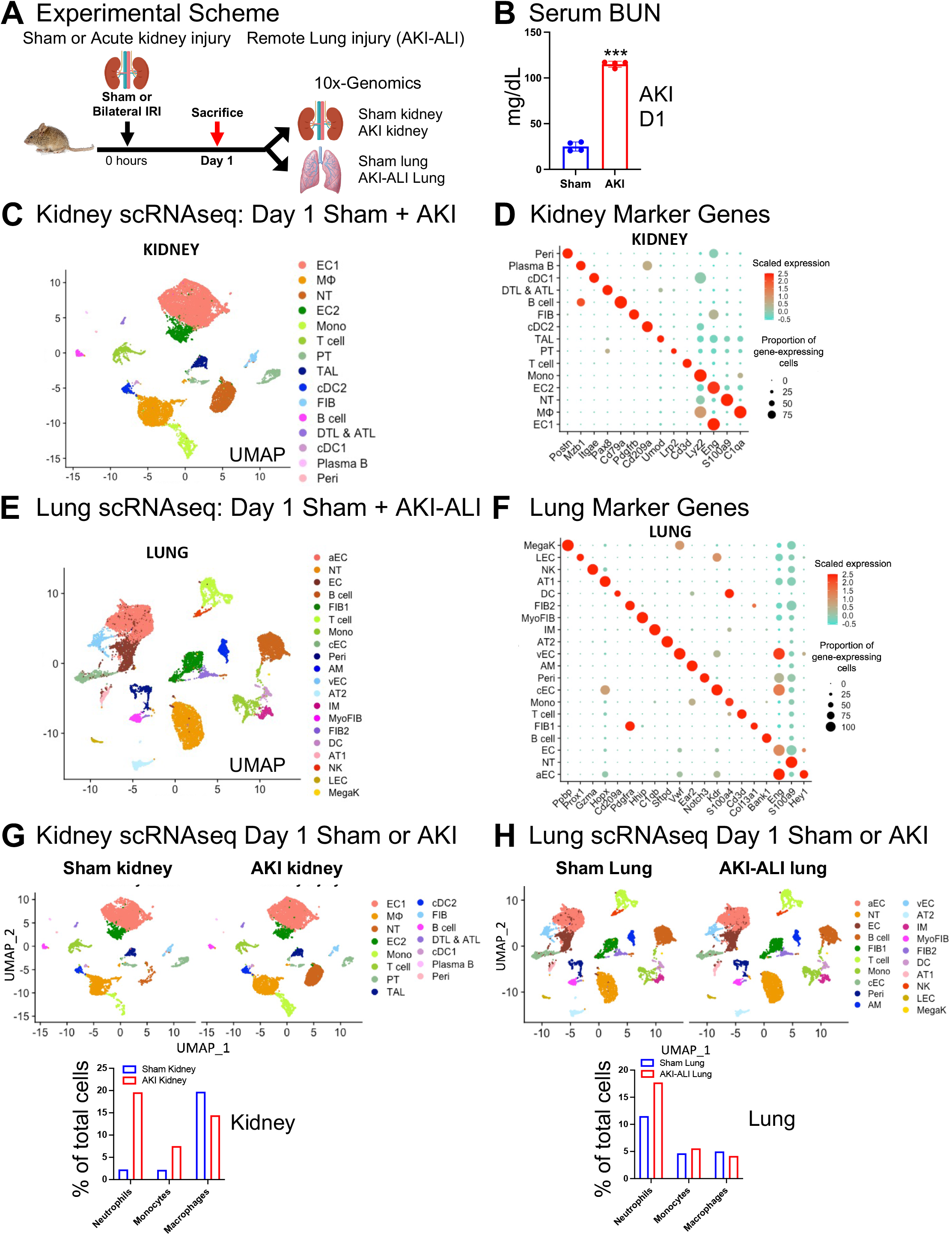
scRNAseq of kidney and lung in setting of AKI-induced ALI (AKI-ALI) (**A.**) Experimental scheme: scRNAseq 10x genomics of kidney and lung Day 1 after sham or AKI, (**B.**) Serum BUN values after Sham or AKI Day 1, (**C.**) Seurat object: combined kidney sham and AKI Day 1 (**D.**) Marker genes used for definition of kidney cell types, (**E.**) Seurat object: combined lung sham and AKI-ALI lung Day 1, (**F.**) Marker genes used for definition of lung cell types, (**G.**) Seurat objects: sham kidney vs. AKI kidney Day 1, quantification of neutrophils, monocytes and macrophages in sham or AKI samples expressed as % of total cells detected by Seurat, (**H.**) Seurat objects: sham lung vs. AKI-ALI lung Day 1, quantification of neutrophils, monocytes and macrophages in sham or AKI samples expressed as % of total cells detected by Seurat. Cell types: *Epithelial:* AT1/2=Type 1/2 lung epithelial cell, PT=Proximal Tubule, TAL=Thick ascending limb, DTL=Descending thin limb, ATL=Ascending-type thin limb, *Endothelial:* EC=Endothelial cell (a=arterial, v=venous, c=capillary), *Stromal:* FIB=Fibroblast, PERI=Pericyte, *Immune cells:* NT=Neutrophil, DC=Dendritic cell, MΦ=Macrophage, AM=Alveolar macrophage, IM=Interstitial macrophage, Mono=Monocyte, MegaK=Megakaryocyte, Plasma B=Plasma B cell, B cell, T cell. n=4 animals per measurement, *** p<0.001.

### Ligand-receptor pairing analysis across organs, linking ligands expressed in the kidney to receptors expressed in the lung

To infer possible cell-cell communication events between kidneys and lungs, we used the machine learning algorithm CellphoneDB^20^ to perform computational Ligand-Receptor (L-R) pairing analysis across organs, with ligands located in the kidney and receptors in the lung (**Figure 3A**). The CellphoneDB Ligand-Receptor interaction database used here is unique in that beyond classical ligand and receptor interactions it also considers other interaction partners that may participate in signaling, such as co-receptors or other receptor-associated proteins. We considered only Ligand-Receptor interactions with the ligand expressed in a kidney cell type and its cognate receptor expressed in a lung cell type. Of note, CellphoneDB calculates an empirical p-value, with higher p-values indicating higher predicted significance of the L-R pairing.

**Figure 3:**
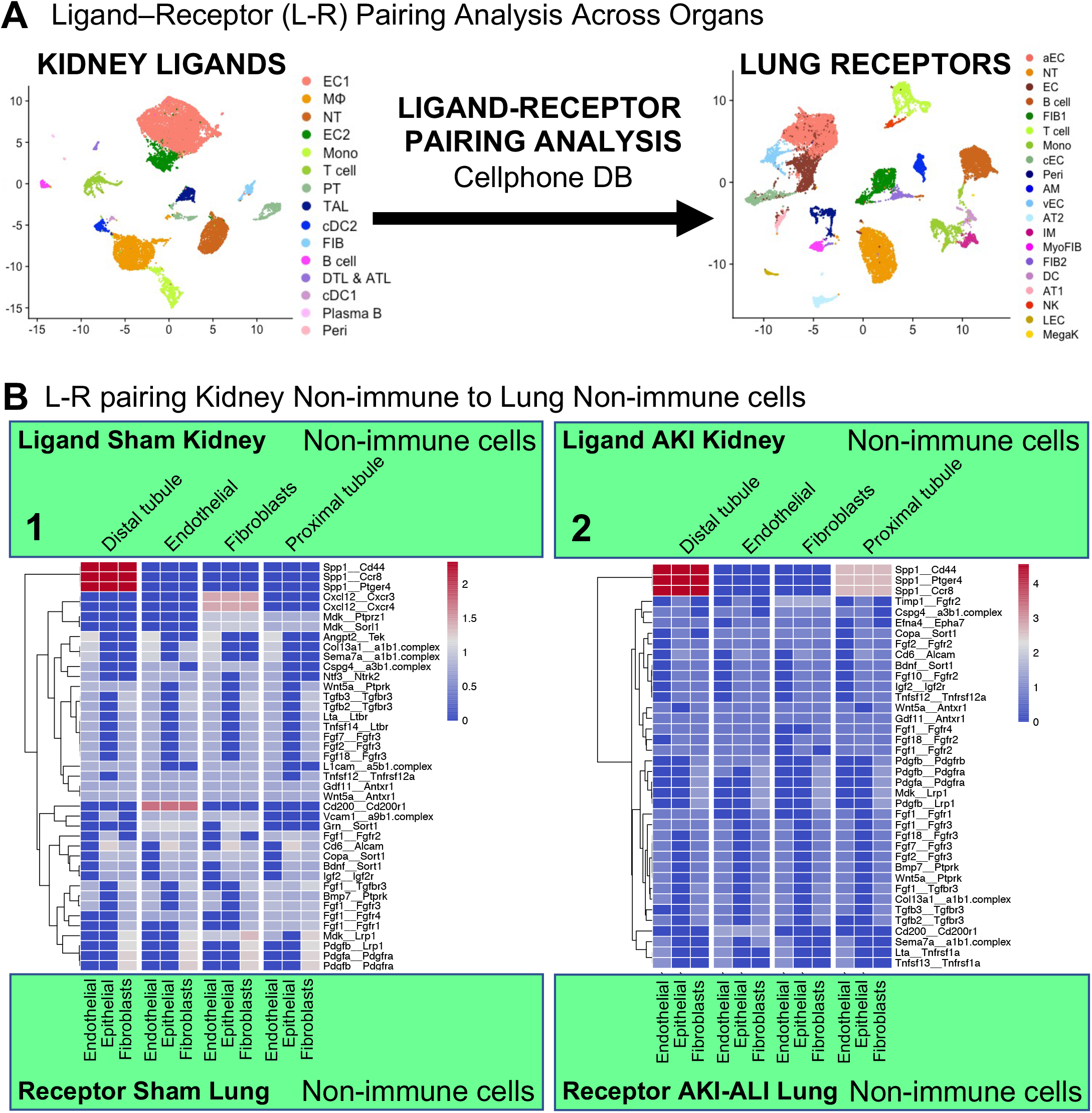
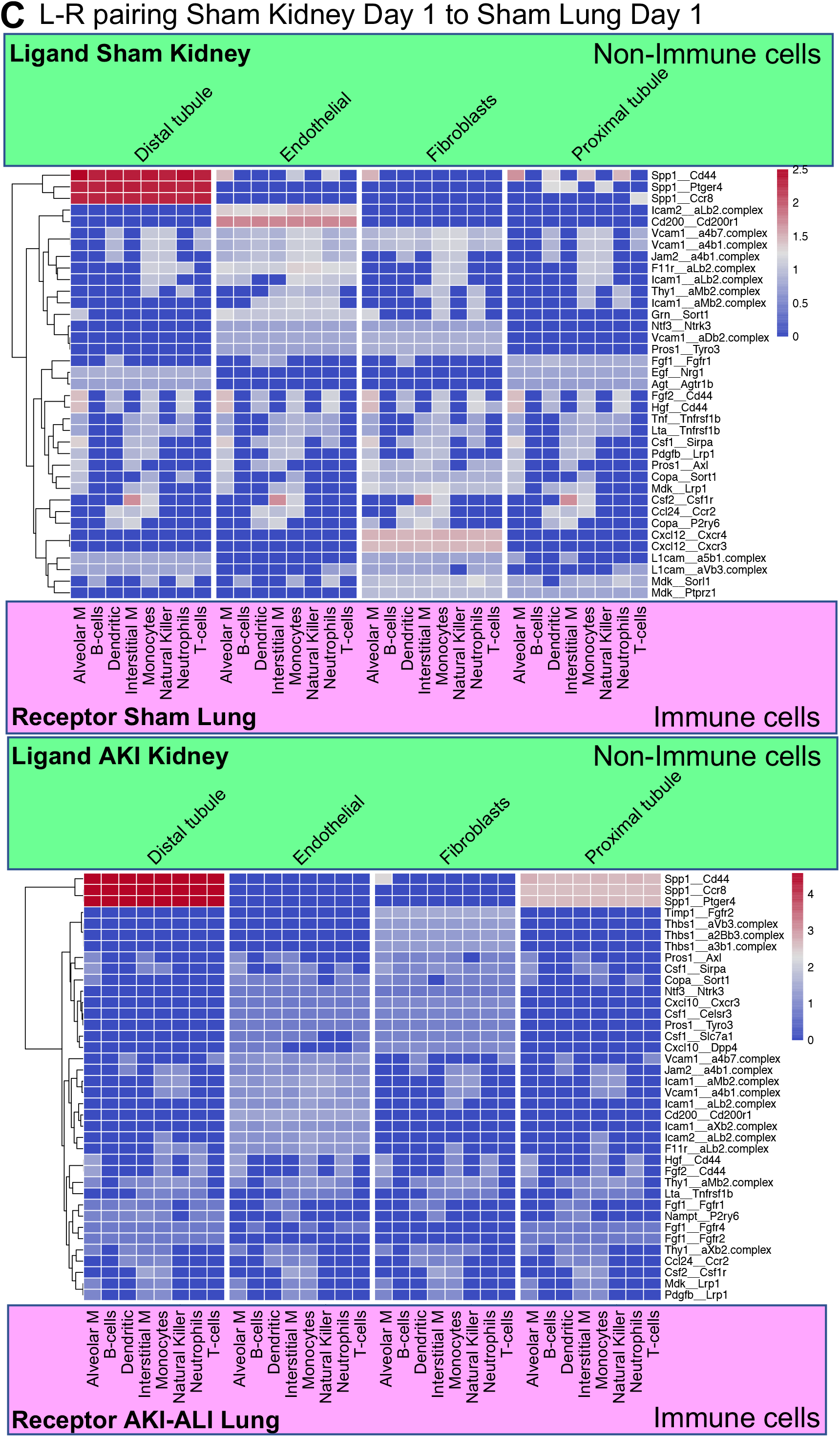
Ligand-receptor pairing analysis across organs, linking ligands expressed in the kidney to receptors expressed in the lung. (**A.**) Experimental scheme: Design of Ligand-Receptor Pairing Analysis, kidney ligands paired to lung receptors using kidney and lung scRNAseq data and CellphoneDB analysis, (**B.**) L-R pairing analysis kidney non-immune cells to lung non-immune cells for Day 1 after sham (panel 1) or AKI (panel 2). p values shown are empirical p values calculated by CellphoneDB, higher values indicate higher L-R pairing significance.

We divided our analysis into L-R pairings from non-immune cells or immune cell types in the kidney to non-immune cells or immune cell types in the lung, either at baseline (sham) or after injury (AKI kidney or AKI-ALI lung). The top scoring L-R pairing detected at baseline and after injury was tubule cell-expressed osteopontin (OPN or secreted-phosphoprotein-1 (SPP1)) pairing with its receptor CD44 in lung non-immune and immune cell types. The significance of these predicted interactions was p > 2.5 at baseline, which significantly increased to p > 4.5 after injury. At baseline, distal tubule cells represented the main source of OPN, but by Day 1 after AKI proximal tubule cells were also predicted to significantly participate in OPN:CD44 signaling (**Figure 3B+C Panels 1 and 2**). CD44 has several other ligands, some of which were also detected in the L-R pairing analysis, such as HGF:CD44 or FGF2:CD44, but these pairings were limited to kidney immune to lung immune cell pairings and of low to moderate significance that did not change with injury (**Supplemental Figure 1A Panel 1 and 2**). For kidney immune cells the top scoring L-R pairings, however with low difference in significance between sham and injured samples (p value >2.5 → 3) included: i.) Kidney neutrophil interleukin-1-beta (IL1b) connecting to adrenoreceptor-beta2 (IL1b:Adrb2) in lung immune or stromal cells, and ii.) kidney T-cell chemokine-ligand-5 (CCL5) connecting to CCR1-5/Ackr4 receptors in lung immune or non-immune cells (**Supplemental Figure 1A+B Panels 1 and 2**). The significance of the IL1b:Adrb2 pairing is unclear, as this pairing is predicted by CellphoneDB based on protein-protein interaction data curated from various experimental approaches^21,22^, but it is not known whether this interaction produces a cellular signal. Whether T-cell released CCL5 or CCL5 in general is involved in AKI-ALI is unknown.

We validated our CellphoneDB analysis using two approaches: <u>First</u>, we used a previously published single-nucleus RNAseq (snRNAseq) dataset that assessed kidney gene expression at early time points after AKI for our L-R pairing analysis^18^. We linked their kidney snRNAseq dataset of 4 and 12 hours after AKI to our scRNAseq data of the remotely injured AKI-ALI lung at Day 1 after AKI. L-R pairing analysis identified OPN:CD44 as a top hit. Kidney distal tubule OPN pairing with CD44 receptor in lung immune and non-immune cell types showed high significance at 4 hours after AKI. At 12 hours after AKI, OPN derived from several kidney cell types including distal tubule cells, endothelial cells, fibroblasts, macrophages, and T cells pairing with CD44 in lung immune or non-immune cells represented the top scoring L-R pairing (**Supplemental Figure 2A+B**). <u>Second</u>, we used a list of all secreted proteins identified in UNIPROT and added a hand-curated list of additional secreted or metalloprotease released proteins of interest and assessed their average expression in all cell types of our AKI kidney scRNAseq dataset. In this analysis, OPN also emerged as the top distal tubule secreted molecule after AKI, and none of the secreted proteins, which were not included in CellPhoneDB, showed a comparable increase (data not shown).

Osteopontin (OPN/SPP-1), initially identified as a regulator of bone biomineralization and remodeling, is an immunoregulatory molecule expressed in a variety of cells, including stromal, epithelial, and immune cells^23–27^. OPN is strongly chemotactic for immune cells, in particular for neutrophils and macrophages, and enhances Th1 inflammation^28–30^. Taking these immunoregulatory features of OPN and our L-R pairing analysis and in silico validation into account, we hypothesized that OPN might be a good candidate AKI-ALI mediator to study in our AKI-ALI model.

### OPN is upregulated in kidneys but not lungs during AKI-induced ALI

To understand the relationship between OPN and the development of AKI-ALI, we assessed expression patterns of OPN mRNA and OPN protein serum levels during the course of AKI. Based on our scRNAseq data, OPN mRNA is already expressed at significant levels in distal tubule and to a lesser degree in proximal tubule cells of sham kidneys (**Figure 4A upper panel**). On Day 1 after AKI OPN expression was most significantly increased in both proximal and distal tubule cells, but now also present at lower levels in other cell types such as endothelial cells, fibroblasts, neutrophils, macrophages, and T cells (**Figure 4A lower panel**). This suggests that tubule-released OPN could potentially act in the induction phase of AKI-ALI, and that at later stages various cellular OPN sources, including immune cells, might contribute to elevated OPN serum levels. We validated these findings by comparing our scRNAseq data with previously published snRNAseq AKI data^18^. Like in our dataset, OPN mRNA expression in sham kidneys was present in both proximal (PTC and New PTC) and distal tubule cells (CNT, DTL-ATL, DCT, Uro). By 4 hours after AKI, OPN mRNA expression began to increase significantly in several cell types and by 12 hours, OPN mRNA expression was very high in proximal and distal tubules, endothelial cells, fibroblasts, neutrophils, macrophages, and T cells, like in our dataset on Day 1 after AKI. OPN expression remained elevated for several weeks in distal tubular compartments (**Figure 4B**). In contrast to the kidney, OPN expression in sham lung was only present in moderate levels in pericytes and to a lesser degree in resident alveolar macrophages. AKI injury did not change the overall low OPN expression levels detected by scRNAseq in the lung on Day 1 after AKI (**Figure 4C upper and lower panel**), suggesting that the kidney but not the lung represents a significant source of OPN after AKI. To better understand dynamic changes in our model, we performed additional AKI-ALI experiments evaluating earlier time points. We subjected C57BL/6 mice to AKI and sacrificed them 1, 2, 4, 6 and 12 hours later (**Experimental Scheme Figure 4D**). In Figures 4+5 we are also incorporating AKI-ALI data from Day 1,3, or 5 after AKI derived from Figure 1 for ease of viewing. Serum BUN and serum creatinine levels were already significantly elevated 1 hour after AKI (**Figure 4E+F**) and serum kidney-injury-molecule-1 (KIM-1, a sensitive marker of kidney injury) was elevated at 12 hours (**Figure 4G**), indicating significant kidney injury. Kidney OPN expression was already significantly elevated 2-4 hours after AKI, reached a peak by 12 hours and remained elevated until at least Day 5. Lung OPN expression, however, remained virtually undetectable or very low over the same time (**Figure 4H**). Serum OPN protein levels closely followed changes in kidney OPN mRNA expression (**Figure 4I**). The early rise in OPN serum levels is likely further enabled by the existing baseline expression of OPN in the sham kidney (**Figure 4A+B**). Elevated OPN levels in AKI-injured mice correlated with the degree of kidney injury (BUN/OPN) (**Figure 4J**).

**Figure 4:**
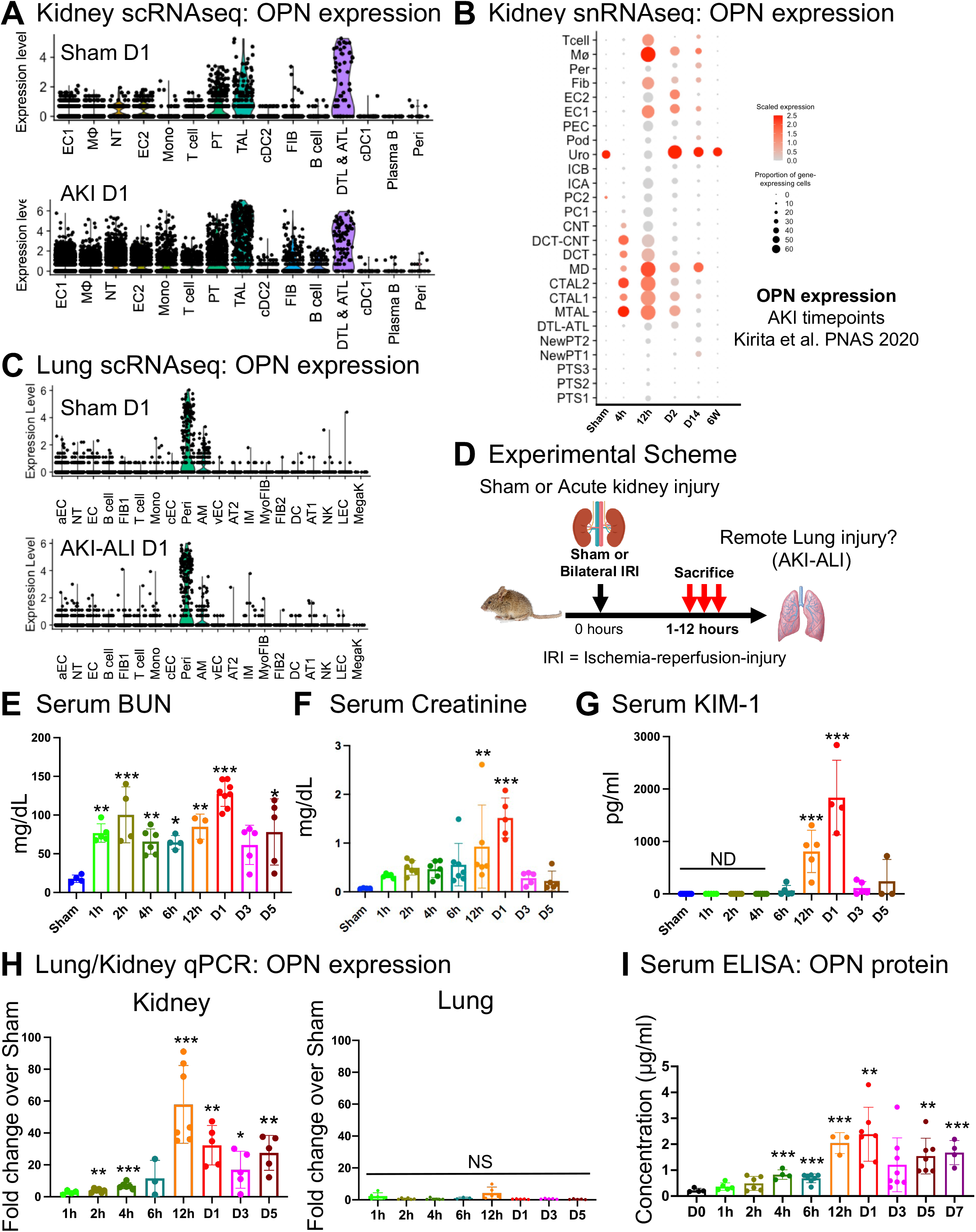
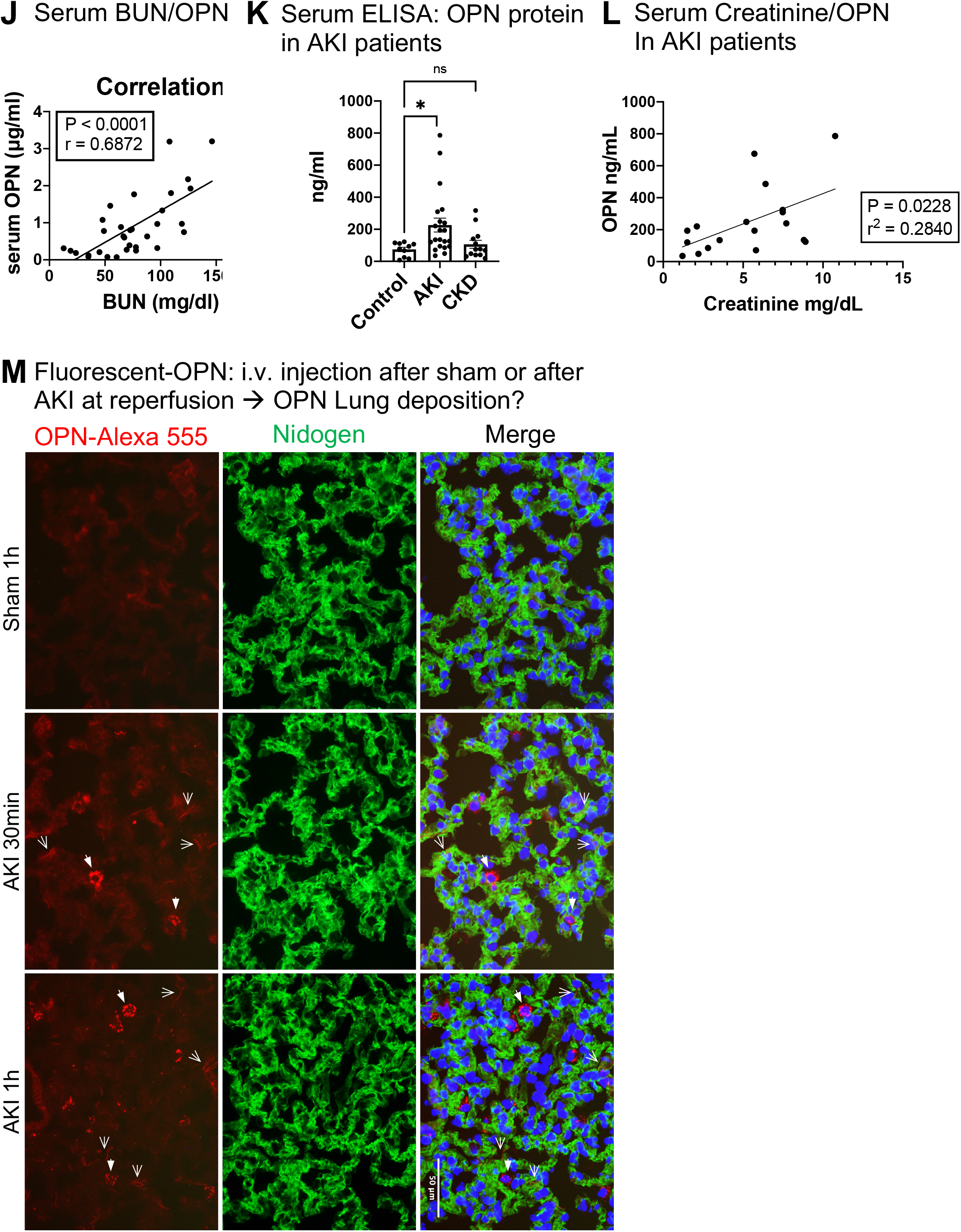
OPN is upregulated in kidneys but not lungs during AKI-induced ALI. (**A.**) Kidney scRNAseq: OPN expression after sham (upper panel) or AKI (lower panel), (**B.**) Kidney snRNAseq (Kirita et al. PNAS 2020): OPN expression after sham or 4hours, 12hours, 2 days, 14 days, and 6 weeks after AKI, (**C.**) Lung scRNAseq: OPN expression after sham (upper panel) or AKI (lower panel), (**D.**) Experimental scheme: AKI → AKI-ALI model. Sham, 1hour, 2hour, 4hour, 12hour after AKI. We integrated measurements from Day 1, 3 and 5 after AKI (Figure 1) in some panels of Figure 4, (**E.**) Serum BUN after Sham or 1-12hours and Day 1,3,5 after AKI, (**F.**) Serum creatinine after Sham or 1-12hours and Day 1,3,5 after AKI, (**G.**) Serum KIM-1 after Sham or 1-12hours and Day 1,3,5 after AKI, (**H.**) Kidney + lung qPCR: OPN expression after Sham or 1-12hours and Day 1,3,5 after AKI, (**I.**) Serum ELISA: OPN protein after Sham or 1-12hours and Day 1,3,5 after AKI, (**J.**) Serum BUN/OPN correlation using BUN and OPN measurements across all time points, **(K.)** Serum ELISA: Human OPN protein levels in patients with AKI compared to healthy controls and patients with chronic kidney disease (CKD), (**L.**) Correlation of OPN serum levels with serum creatinine concentration in patients with AKI. (**M.**) Fluorescent OPN injection: OPN Alexa 555 (red) in lung tissue at 1 hour after Sham or 30min-1hours after AKI (injection at reperfusion or similar time in sham). The extracellular matrix component Nidogen (green) was used to localize extracellular/interstitial space. DAPI stain (blue) was used to visualize nuclei. Open arrows = OPN Alexa 555 accumulation in lung interstitium, closed arrows = OPN Alexa 555 uptake into lung cells with perinuclear localization. n=3-8 animals per measurement, *** p<0.001, ** p<0.01, * p<0.05. NS=not significant. ND=not detected

**Figure 5:**
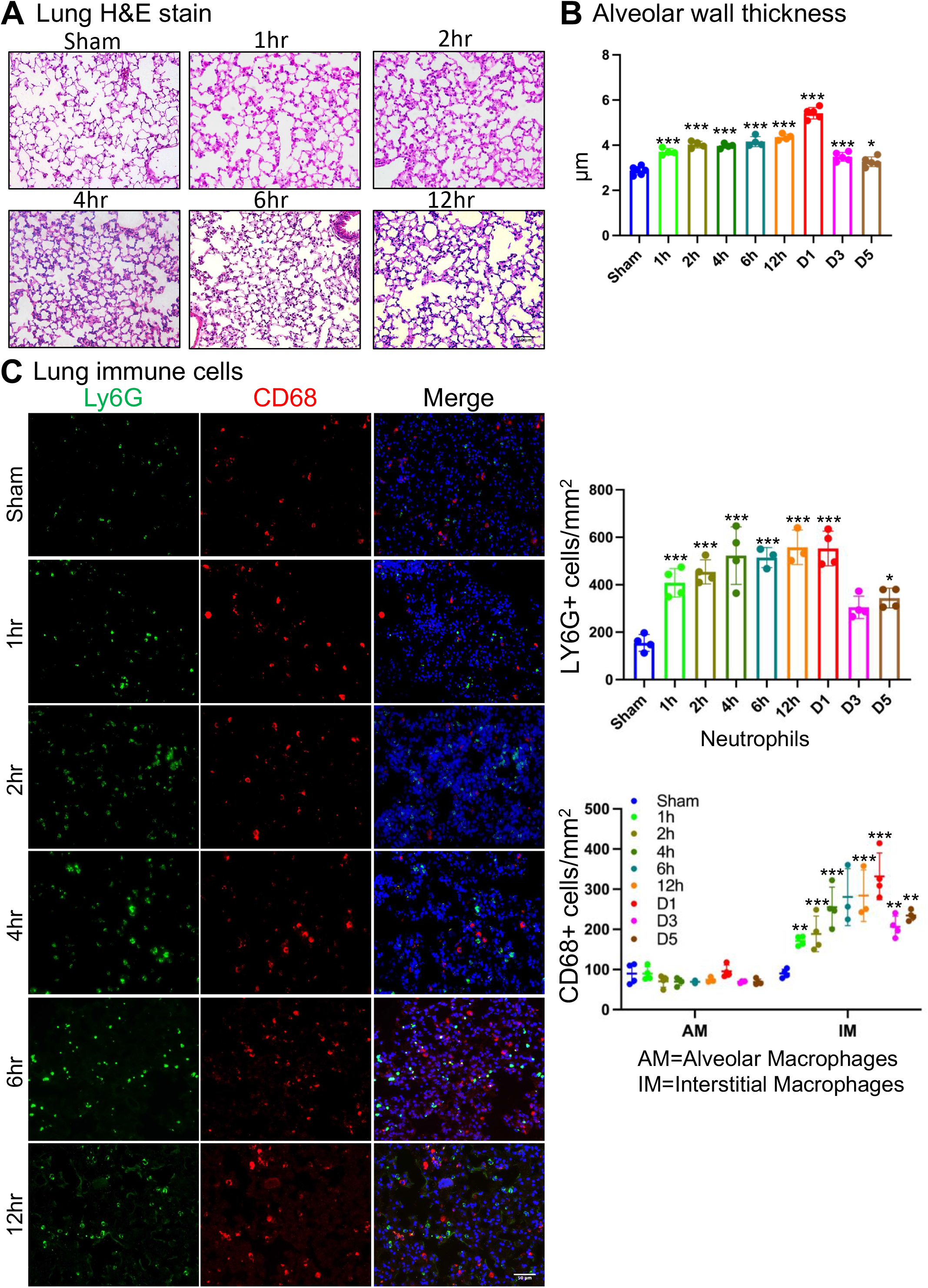
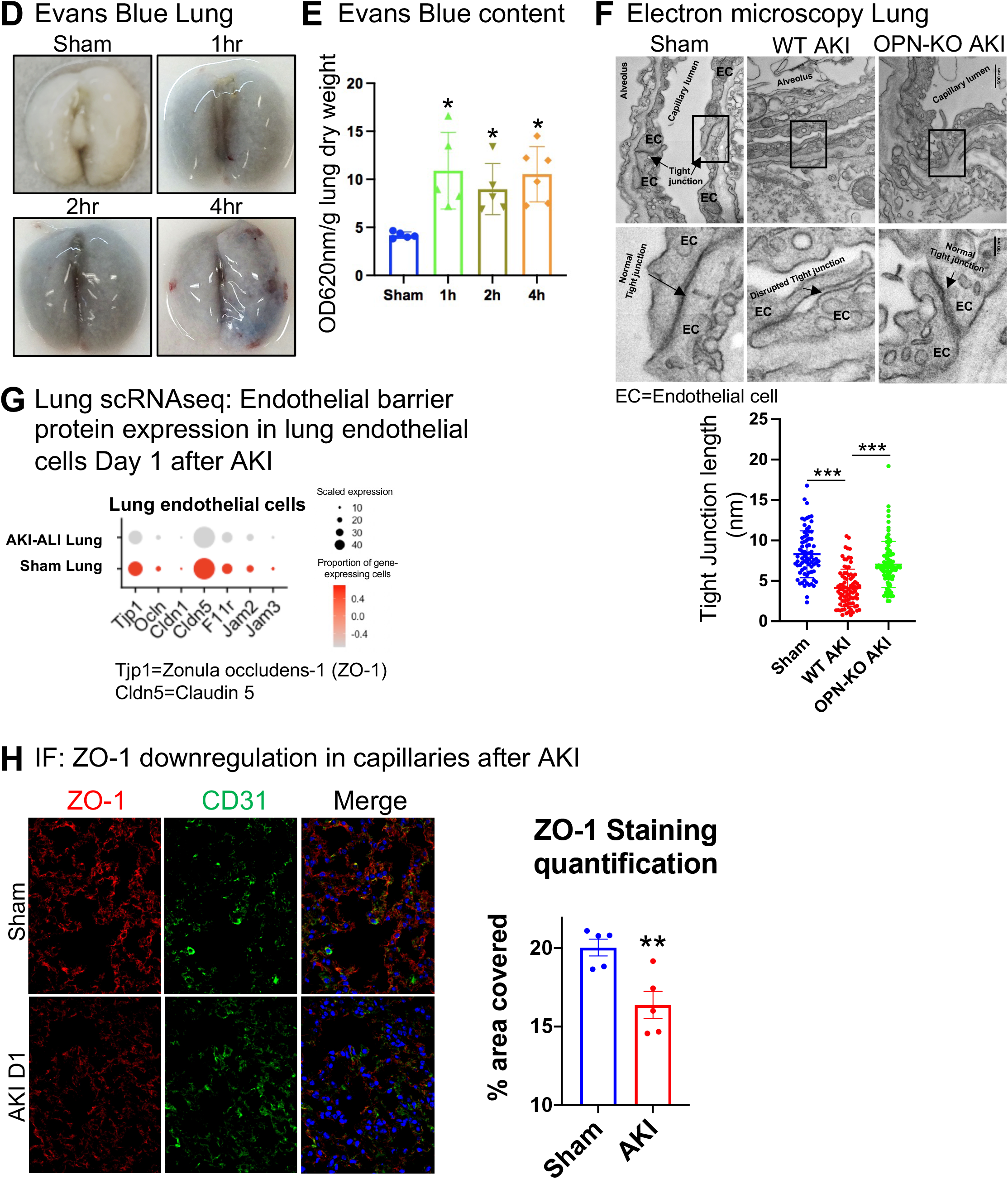
AKI triggers lung endothelial barrier dysfunction and immune cell accumulation very early after AKI. (**A.**) Lung H&E stain after Sham or 1-12hours after AKI, (**B.**) Alveolar thickness measurements after Sham or 1-12hours and Day 1,3,5 after AKI, (**C.**) Lung neutrophils (Ly6G^+^, green), alveolar (CD68^high^, large, red) and interstitial macrophages (CD68^low^, small, red) and quantification after Sham or AKI Day 1,3,5 after AKI. DAPI stain (blue) was used to visualize nulei, (**D.**) Lung Evans Blue leakage after sham or 1-4hr after AKI, (**E.**) Lung Evans Blue quantification (OD620nm/g lung dry weight) after sham or 1-4hr after AKI, (**F.**) Lung electron microscopy and quantification of endothelial tight junction length (nm) at Day 1 in sham, or Day 1 after AKI in *wt* control or OPN-global-KO mice, (**G.**) Lung scRNAseq: Expression of endothelial barrier proteins in lung endothelial cells Day 1 after sham or AKI; Tjp1=Zonula-occludens-1 (ZO-1), Cdnl5=claudin-5, (**H.**) Lung ZO-1 protein expression (IF): ZO-1 (red), CD31 (green, marking endothelial cells), DAPI (blue, marking nucleus) and quantification Day 1 after sham or AKI, (**I.**) Lung qPCR: ZO-1 expression Day 1 after sham or AKI,. n=3-6animals per measurement, *** p<0.001, ** p<0.01, * p<0.05.

Serum OPN protein levels have been studied as a biomarker of severity of disease in patients with multiorgan failure, often including AKI and ALI^31,32^. Whether serum OPN levels are also elevated in patients with AKI prior to development of multiorgan failure is not known. We examined OPN serum levels in patients with AKI that did not have multiorgan failure and were not critically ill. Serum OPN levels were significantly elevated in AKI patients, as compared to healthy controls or patients with chronic kidney disease (CKD) (**Figure 4K**) and positively correlated with reduced kidney function as determined by serum creatinine measurements (**Figure L**). These findings suggest that AKI in humans could result in release of significant amounts of OPN into the circulation and raise the possibility that circulating OPN may promote damage at remote sites also in humans.

We hypothesized that for OPN to act on lung and cause AKI-ALI it might need to gain access to the lung tissue compartment from the circulation. Alternatively, it could act on cells directly accessible from the circulation, such as endothelial cells or circulating immune cells. We thus examined whether OPN can gain access to lung tissue from the circulation in the setting of AKI in mice. We injected fluorescently labeled OPN after sham operation or after AKI at the time of reperfusion and sacrificed animals 30 minutes and 1 hour after injection. We co-stained for nidogen, a marker of the extracellular matrix/interstitial space. Sham operated animals did not accumulate any significant amount of fluorescently marked OPN in the lung within 1 hour of injection, whereas animals with AKI showed OPN accumulation in lung interstitium (open arrows) and uptake into lung cells with perinuclear localization (closed arrows) at 30 minutes-1 hour after injection (**Figure 4M**). Taken together, our results indicate that kidney rather than lung represents the source of circulating OPN protein after AKI and that circulating OPN can principally gain access to the lung early after AKI and thus might mediate the induction of remote lung injury. Access to the lung was marginal in Sham animals, leading us to the conclusion that another AKI-released factor must cooperate with OPN to allow OPN access to lung tissue.

### AKI triggers lung endothelial barrier dysfunction and immune cell accumulation very early after injury

Consistent with the timing of elevated OPN serum levels after AKI, lungs already showed significant signs of injury with interstitial expansion and increased cellularity by 1 hour after AKI which worsened progressively over the subsequent 12 hours (**Figure 5A, also compare to Figure 1D**). Alveolar thickness increased significantly by 1 hour (>1.3fold increase over sham), worsened progressively over the ensuing 12 hours (>1.5fold increase over sham) and reached a peak at day 1 after AKI (>1.85fold increase over sham) (**Figure 5B**). Consistent with these findings, accumulation of neutrophils and macrophages showed a similar trend to the timeline of changes in kidney OPN mRNA expression and serum OPN levels (**Figure 5C**).

Endothelial homeostasis is disrupted in acute lung injury or its most severe form, acute-respiratory-distress-syndrome (ARDS)^33^, which often occurs in response to direct lung injury, such as by COVID-19 lung infection^34^, or as a secondary organ complication of sepsis^35–37^. Severe acute lung injury/ARDS is characterized by diffuse endothelial injury, intense activation of the coagulation system and increased capillary permeability^34,38^. Increased vascular barrier permeability has also been reported in animal models of AKI^15,39–41^. We thus set out to examine whether endothelial barrier disruption, a hallmark of lung injury, also occurs following AKI in our model. Vascular leakage, as evidenced by Evans blue dye uptake, was already detectable at a level twice as high as sham controls at 1 hour after AKI, which was maintained for at least 4 hours (**Figure 5D+E**). Electron microscopy also showed that, compared to sham controls, endothelial tight junctions in AKI-injured C57BL/6 animals were shortened in length or disrupted in Day 1 AKI-ALI lungs. By contrast, AKI-injured OPN-global-KO mice displayed normal endothelial tight junctions, comparable to sham operated *wt* animals (**Figure 5F**). We next assessed lung endothelial cells. Using our scRNAseq data, we found that the expression levels of two endothelial barrier proteins, zonula occludens-1 (ZO-1, Tjp1) and claudin-5 (Cldn5), were strongly downregulated in AKI-ALI samples as compared to sham lungs (**Figure 5G)**. We validated our finding on the protein level for ZO-1. ZO-1 staining was heavily present in the capillary network of sham lung, but severely diminished in the capillary network of the AKI-ALI lung. CD31 (green) was used as a co-stain to identify endothelial cells (**Figure 5I**). Other cell types in the lung, in particular stromal cells, such as fibroblasts and lung epithelial cells (AT1/2) also expressed ZO-1 or claudin-5 at low levels and these expression levels did not significantly change in the AKI-ALI lung (data not shown). These results suggest that endothelial barrier dysfunction and vascular leakage represent early events in the induction of AKI-ALI remote lung injury in our model.

### Pharmacological or genetic inhibition of OPN protects from ALI after AKI

To assess whether OPN is necessary for the development of AKI-ALI, we subjected C57BL/6 mice to bilateral ischemia-reperfusion-injury and injected the animals with either control IgG or anti-OPN neutralizing antibody. Analogous experiments were performed in OPN-global-knockout or *wt* mice without antibody treatments (**Experimental Scheme Figure 6A**). Anti-OPN-antibody treated mice or OPN-global-knockout mice experienced the same degree of kidney injury on day 1 as controls, as evidenced by comparable elevations in serum BUN and KIM-1 levels (**Figure 6B+C**). However, lung injury was significantly ameliorated after treatment with anti-OPN-antibody and in OPN-global-knockout mice (**Figure 6D+E**). While anti-OPN-antibody strongly reduced lung neutrophil and interstitial macrophage accumulation after AKI, OPN-global-KO prevented it almost completely (**Figure 6F**). Anti-OPN-antibody-treated animals also showed strongly reduced vascular leakage and improved lung function as compared to controls on day 1 after AKI (**Figure 6G-I**). Taken together, our results identify circulating OPN as a critical and necessary regulator of lung endothelial barrier permeability, lung immune cell accumulation and functional impairment in AKI-ALI.

**Figure 6:**
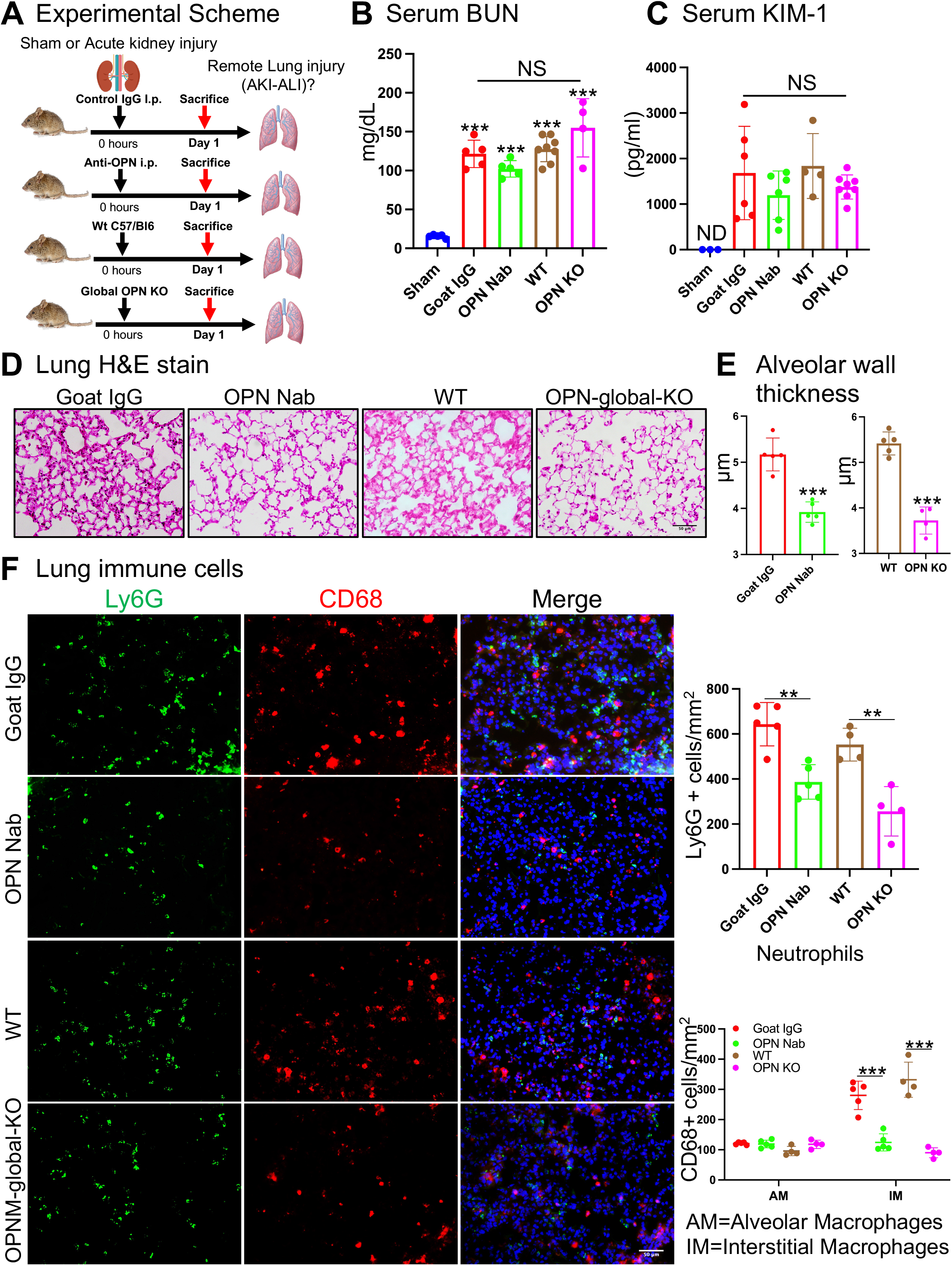
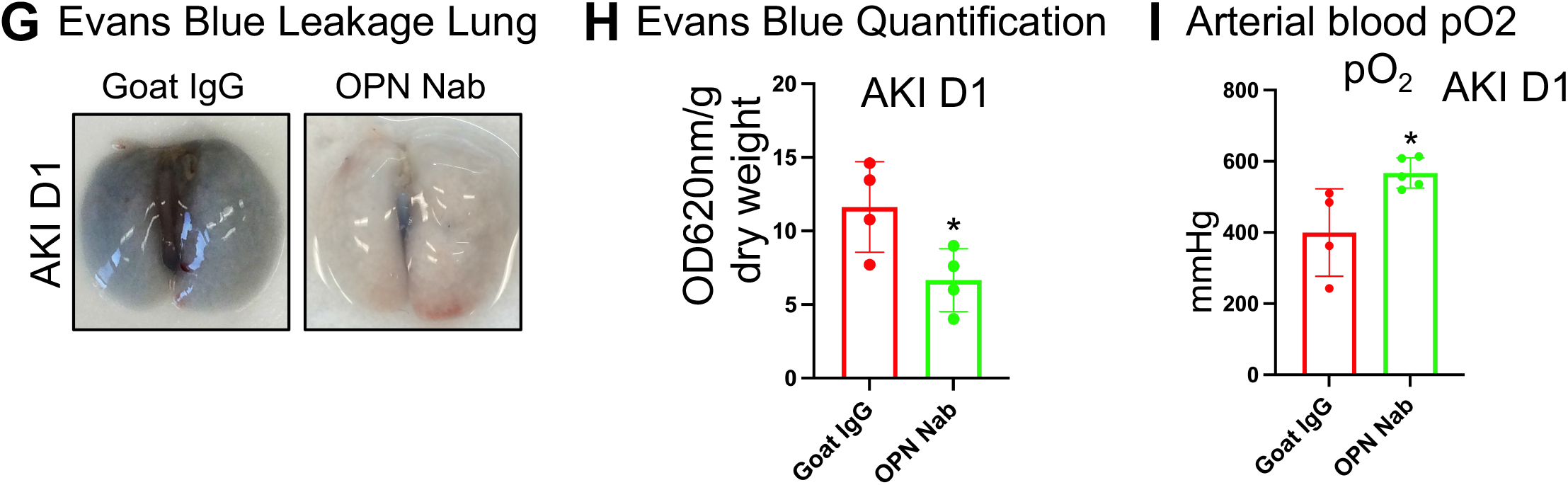
Pharmacological or genetic inhibition of OPN protects from ALI after AKI. (**A.**) Experimental scheme: AKI → AKI-ALI model Day 1 after AKI. In the following *wt* mice injected with control goat IgG are compared to OPN-neutralizing-antibody (OPN Nab) injected mice, and OPN-global-KO animals (OPN KO) are compared to *wt* controls, (**B.**) Serum BUN values after Sham or AKI Day 1, (**C.**) Serum creatinine values after Sham or AKI Day 1, (**D.**) Lung H&E stain after Sham or AKI Day 1, (**E.**) Alveolar thickness measurements after Sham or AKI Day 1, (**F.**) Lung neutrophils (Ly6G^+^, green), alveolar (CD68^high^, large, red) and interstitial macrophages (CD68^low^, small, red) and quantification after Sham or AKI Day 1. DAPI stain (blue) was used to visualize nuclei, (**G.**) Lung Evans Blue leakage Day 1 after sham or AKI, (**H.**) Lung Evans Blue quantification (OD620nm/g lung dry weight) Day 1 after sham or AKI, (**I.**) Arterial blood oxygen partial pressure after Sham or AKI Day 1. n=3-8 animals per measurement, *** p<0.001, ** p<0.01, * p<0.05. NS=not significant.

### Circulating OPN is sufficient to induce ALI after AKI

To determine whether OPN suffices to mediate AKI-ALI, we tested the effect of OPN injection in the context of mild AKI (reduced ischemia time), an experimental condition where AKI-ALI is not detectable at 6 hours after AKI (**Experimental Scheme Figure 7A**). As expected, serum OPN levels were significantly lower in mild AKI than severe AKI (**Figure 7B**). Serum BUN levels were significantly elevated at 6 hours after mild AKI compared to sham controls, but lower compared to severe AKI (**Figure 7C).** Intravenous injection of OPN protein into mild AKI animals at the time of kidney reperfusion at quantities that would mimic serum concentrations at 6 hours after severe AKI (see Figure 4I) triggered AKI-ALI that was now comparable to severe AKI-ALI (**Figure 7E+F**). We also performed OPN injections into uninjured mice but were unable to detect increases in alveolar thickness or inflammatory changes in the lung (**Supplemental Figure 3**), suggesting that a second event or mediator induced by AKI is required for OPN to successfully induce AKI-ALI. Therefore, we conclude that OPN is sufficient to induce AKI-ALI only in the context of AKI.

**Figure 7:**
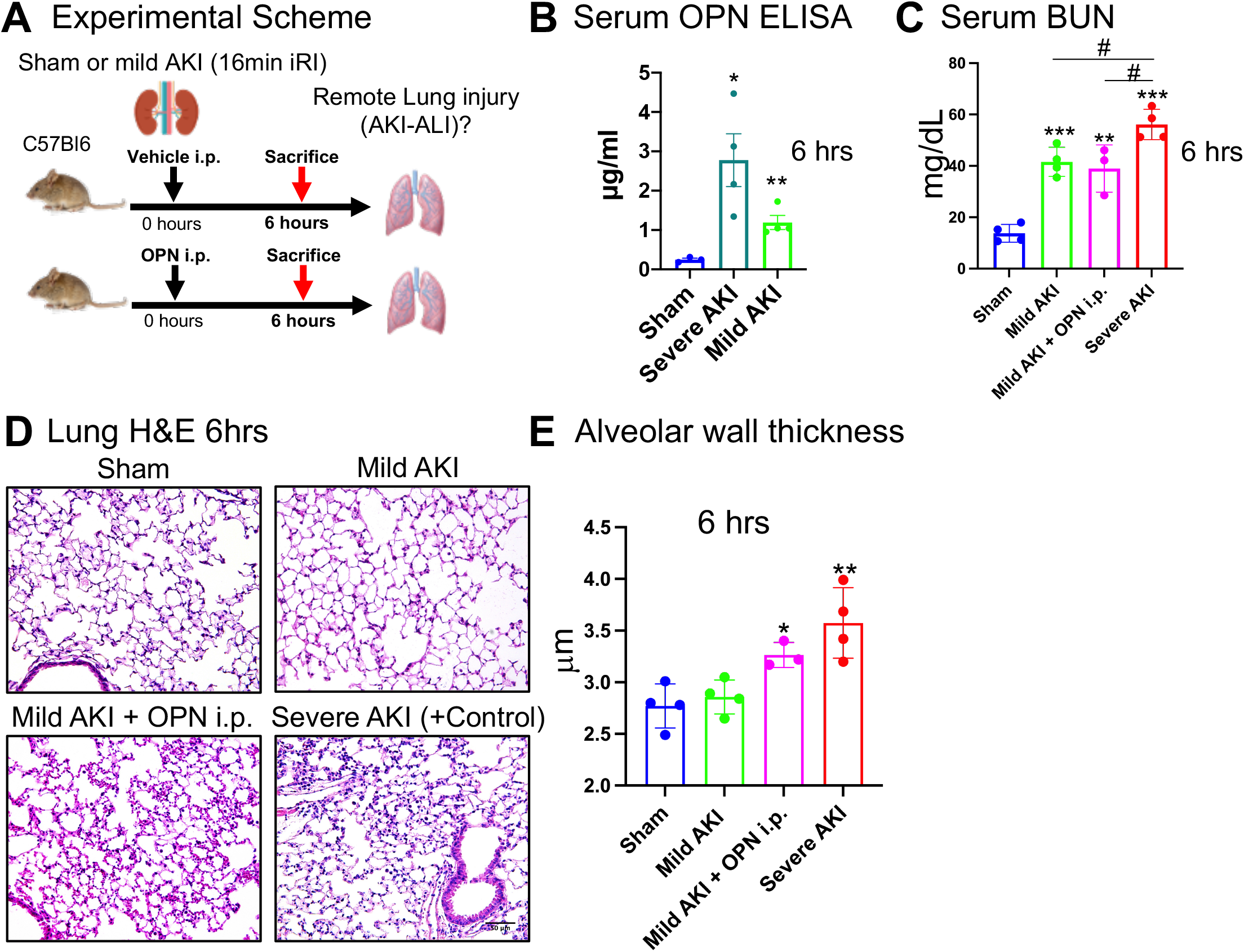
Circulating OPN is sufficient to induce ALI after AKI. (**A.**) Experimental scheme: AKI → AKI-ALI model evaluated 6 hours after mild AKI (reduced ischemia time). Mice with mild AKI are injected with vehicle control or OPN protein, severe AKI is used as a control, (**B.**) Serum ELISA: OPN protein 6 hours after sham, severe AKI (positive control) or mild AKI, (**C.**) Serum BUN 6 hours after sham, mild AKI +/− OPN injection or severe AKI (positive control), (**D.**) Lung H&E stain 6 hours after sham, mild AKI +/− OPN injection or severe AKI (positive control), (**E.**) Alveolar thickness measurements 6 hours after sham, mild AKI +/− OPN injection or severe AKI (positive control). n=3-4 animals per measurement, *** p<0.001, ** p<0.01, * p<0.05, # p<0.05.

### Circulating OPN relevant for induction of AKI-ALI is released from the injured kidney

Upregulated OPN expression and serum levels have been found in the context of a number of organ injuries^31,32, 42–48^, allowing for the possibility that OPN sources in the body other than the kidney might be relevant in AKI-ALI. Conversely, it has never been conclusively shown that a proposed AKI-ALI mediator is released directly from the injured kidney. To address this question, we performed kidney transplantation experiments, where we transplanted either ischemic *wt* kidneys or ischemic OPN-global-knockout kidneys into syngeneic C57BL/6 mice (**Experimental Scheme Figure 8A**). KIM-1 kidney mRNA expression and tubular injury scores document that transplanted OPN-global-knockout kidneys show kidney injury parameters comparable to those in transplanted *wt* kidneys (**Figure 8B+C**). Serum BUN and creatinine measurements would be unaffected in our transplantation model, given that two intact kidneys are present in the mouse as well. OPN serum levels were elevated above sham levels in mice that received *wt* kidneys, but not in mice that were transplanted OPN-KO-renal grafts. Of note, OPN serum levels are lower with only one injured transplant kidney, as compared to the bilateral IRI model, where two kidneys are injured (**Figure 8D** and see Figure 4). Lung injury developed in mice that received *wt* kidneys but not in those that received OPN-KO kidney grafts (**Figure 8E-I**). These results conclusively identify the kidneys as the source of OPN during the development of ALI after AKI.

**Figure 8:**
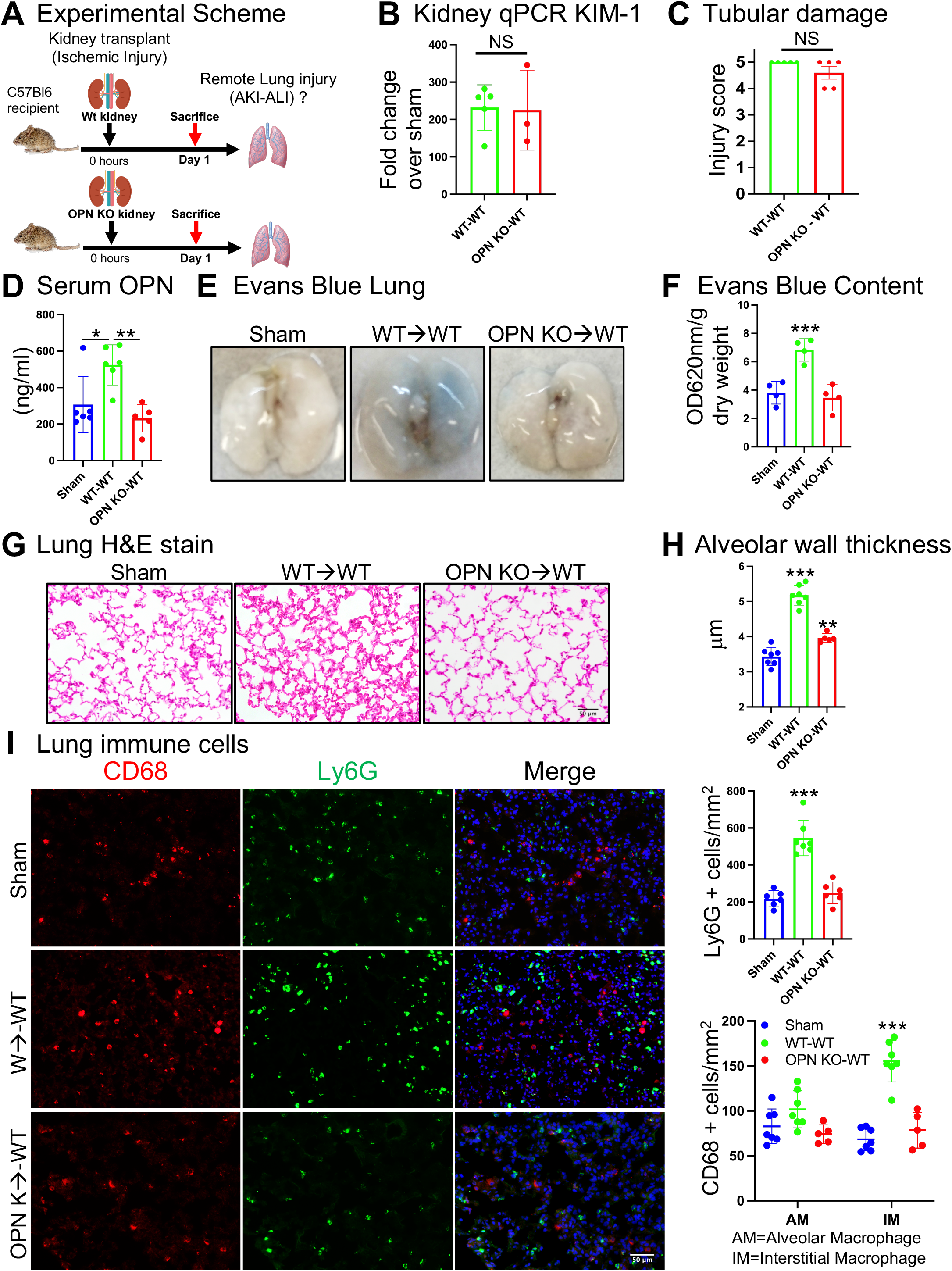
Circulating OPN relevant for induction of AKI-ALI is released from the injured kidney. (**A.**) Experimental scheme: Ischemic kidney transplantation → AKI-ALI at Day 1 after AKI. C57BL/6 *wt* kidneys or OPN-global-KO kidneys were transplanted into C57BL/6 *wt* mice (wt-wt = wt kidney-to-C57Bl/6 transplant and OPN KO-wt = OPN KO kidney-to-C57BL/6 transplant), (**B.**) Kidney qPCR: KIM-1 expression in wt-wt and OPN KO-wt Day 1 after transplant, (**C.**) Tubular injury score of transplanted wt or OPN-global-KO kidney Day 1 after transplant, (**D.**) Serum ELISA: OPN protein in wt-wt and OPN KO-wt Day 1 after transplant, (**E.**) Lung Evans Blue leakage in lungs of *wt* or OPN-global-KO kidney transplanted mice Day 1 after transplant, (**F.**) Lung Evans Blue quantification (OD620nm/g lung dry weight) in lungs of *wt* or OPN-global-KO kidney transplanted mice Day 1 after transplant, (**G.**) Lung H&E stain of *wt* or OPN-global-KO kidney transplanted mice Day 1 after transplant, (**H.**) Alveolar thickness measurements of *wt* or OPN-global-KO kidney transplanted mice Day 1 after transplant, (**I.**) Lung neutrophils (Ly6G^+^, green), alveolar (CD68^high^, large, red) and interstitial macrophages (CD68^low^, small, red) and quantification in lungs of *wt* or OPN-global-KO kidney transplanted mice Day 1 after transplant. DAPI stain (blue) was used to visualize nuclei. n=4-7 animals per measurement, *** p<0.001, ** p<0.01, * p<0.05.

## Discussion

Our work using kidney transplantation provides the first conclusive evidence *in vivo* that a kidney-released circulating mediator is mechanistically involved in secondary organ failure after kidney injury. Neutralization of circulating OPN or OPN-global-KO prevented AKI-ALI and respiratory failure, and ischemic kidneys from OPN-global-KO mice transplanted into *wt* mice failed to raise serum OPN levels and to induce AKI-ALI, as compared to transplantation of ischemic *wt* kidneys into *wt* mice. We show that distal- and proximal tubule-released OPN is poised to act as a key mediator of lung vascular leakage and inflammation early in the lung’s response to kidney injury. The fast response is due to OPN’s baseline expression and quick upregulation, causing lung vascular leakage and immune cell accumulation within an hour, ultimately resulting in functional impairment with reduced arterial oxygenation. This finding has implications beyond acute kidney injury, as tissue injury due to various causes can raise OPN serum levels in humans, including but not limited to direct lung injury by COVID-19, bacterial pneumonia^49,50^, by sepsis from various causes^32,51,52^, or cardiac injury^47,51^. OPN has also been described as a marker of kidney injury and declining kidney function ^e.g.53,54^, but not much is known about the mechanisms involved. All these conditions are highly associated with multiorgan failure, in particular AKI and ALI or its most severe form acute respiratory distress syndrome (ARDS). This suggests that OPN upregulation represents a conserved response to tissue injury in different organs and might indeed link circulating OPN to secondary organ failure in general, as is suggested by various studies of serum OPN as a biomarker of multiorgan failure mortality in humans^32,47,51^. Consistent with this, one other study in mice connected OPN to lung injury after intestinal ischemia^55^. However, in contrast to our study, OPN in this model was found upregulated in both the ischemic gut and the remotely injured lung, not allowing the investigators to identify the source of serum OPN elevations or conclusively determine that circulating OPN was indeed involved in inducing lung injury. Broadening the putative relevance of our study is the fact that remote organ effects after AKI can also be detected in a number of other organs, including the heart and brain^13^.OPN serum levels are elevated in the systemic inflammatory response syndrome (SIRS) and sepsis^52^ and predict mortality in patients with sepsis and multiorgan failure^31,32^.

One published study of patients with multiorgan failure allows to link OPN serum levels in humans to respiratory outcomes. A study of 159 intensive care unit admissions (70% with sepsis, 14% cardiopulmonary disease, 10% liver disease, 21% other diseases, 74% artificial ventilation, many with AKI) shows that elevated OPN serum levels positively correlate with reduced kidney function measured by creatinine (r^2^=0.363, p<0.001) or glomerular filtration rate (r^2^=-0.39, p=0.001) as well as the presence of multiorgan failure (determined by Sequential-Organ-Failure-Assessment (SOFA) score) (r^2^=0.512, p=0.002) and decreased survival (r^2^=-0.342, p=0.017). Consistent with our observations in mice, elevated OPN serum levels in these patients are also positively correlated with respiratory failure and increased need for artificial ventilation (r^2^=0.212, p=0.032)^51^. This data suggests that OPN might have similar roles in AKI-ALI with respiratory failure in humans as we detect in mice.

A study of critically ill patients with multiorgan failure including AKI requiring dialysis found that OPN levels in this cohort were significantly elevated when compared with critically ill controls without AKI^47^, possibly indicating that the degree of kidney injury correlates with the degree of serum OPN elevation. Our results show that serum OPN is indeed elevated in cases of human AKI (no multiorgan failure or sepsis) and correlates with the degree of kidney injury, strengthening the link between elevated OPN serum levels and kidney injury in humans. Our results allow the possibility that elevated serum OPN levels in critically ill patients with AKI are not only associated with respiratory failure^51^, but may indeed be causative. Yet, our findings of OPN serum elevations in AKI patients will need to be followed up with a prospective study that conclusively connects OPN serum levels to respiratory outcomes in humans. Currently there are to our knowledge no already existing human sample collections available to assess this connection.

Our studies also for the first time identify circulating OPN as a key regulator of endothelial barrier permeability *in vivo* in AKI-ALI. A previous study associated OPN with endothelial barrier permeability in pulmonary-vein-endothelial-cells (PUVECs) in primary culture. In this report, OPN expression was increased, and ZO-1 and claudin-5 expression levels were decreased in PUVECs in septic rats or in LPS-treated PUVECs isolated from these rats. Upregulation of connexin43 in PUVECs stimulated OPN expression and downregulated ZO-1 and claudin-5, increasing vascular permeability. OPN knockdown blocked this effect^56,57^. These data are consistent with our findings of ZO-1 and claudin-5 downregulation in single-cell expression data of lung endothelial cells (Figure 5G) and downregulation of ZO-1 protein in immunofluorescence stains of AKI-ALI lung tissue (Figure 5H). Also, principally consistent with our findings, OPN was found upregulated by the host and tumor in a mouse model of malignant pleural effusions where OPN neutralization reduced leakage of fluid into the pleural space^58,59^.

Tumor necrosis factor (TNF) and its family member Lymphotoxin A (Lta), were also found in our L-R pairing analysis, adding additional validity to our strategy and findings. Circulating TNF is thought to participate in the induction of lung cell death via its receptor TNFR1, as neutralization of TNF with etanercept in AKI-injured *wt* mice or use of global TNFR1 knockout mice reduced lung cell apoptosis after AKI^17^. We identified various moderate to low significance pairings of TNF with its cognate receptors, in particular TNF:TNFR1, and other known interacting proteins (**Supplemental Figure 2**). These TNF pairings were primarily identified between kidney immune cells and lung stromal or immune cells when pairing sham kidney with sham lung, or at somewhat higher significance when pairing early AKI kidney (4 hours and 12 hours) with Day 1 AKI-ALI lung. Yet, most of these pairings lost significance when Day 1 AKI kidney was paired with Day 1 AKI-ALI lung (**Figure 3D+E panels 1+2 and Supplemental Figure 2**). This suggests that TNF plays a role early in the development of AKI-ALI, consistent with the role of TNF as a Type 1 inflammatory cytokine known to act early in establishing inflammation, and this is also consistent with the referenced study of TNF inhibition/TNFR1 deficiency in AKI-ALI^17^. Of note, TNF is made as a pro-form that is cleaved on the cell surface to release soluble TNF^60^. We also identified moderate to low significance L-R pairing of kidney TNF with its lung receptor TNFR2 (TNFRSF1b) in immune cells but not stromal cells; however, the significance of these pairings did not increase with injury (**Figure 3B+C+D and Supplemental Figure 2**). TNFR2 is almost exclusively expressed in immune cells, and its signaling requires cell-cell contact, allowing interaction of transmembrane pro-TNF (which is detected in its mRNA form by our scRNAseq experiment) expressed in a signaling cell with TNFR2 in an adjacent cell. This interaction is thus likely not occurring between kidney-released circulating soluble TNF and TNFR2 in the lung but could principally occur if an immune cell that is primed in the injured kidney to express pro-TNF were to travel to the lung. Yet to date there is no evidence for this occurring in AKI-ALI. Similar to TNF, IL6 was linked to AKI-ALI using IL6 neutralization and IL6 global knockout mice^61^. IL6 and its receptor, however, were not identified by our analysis.

We did not find significant lung changes after bilateral nephrectomy. Others have however demonstrated changes in gene expression profiles and lung inflammation after nephrectomy^62,63^, and the elimination of AKI-induced cytokines can indeed be delayed by nephrectomy^64^. In our experiments we can detect small changes in serum OPN even after sham surgeries, without kidney injury or nephrectomy, possibly induced by the relatively small tissue injury induced by sham surgery (data not shown). It is thus possible that the detection of lung changes after nephrectomy depends at least in part on the extent of tissue injury caused by the surgeon. This might explain some of the discrepancies between our work and the work of others.

Osteopontin (OPN), also known as secreted phosphoprotein 1 (SPP-1), early T lymphocyte activation-1 (Eta-1) or uroprotein^25,26,30^ is a member of the small integrin-binding ligand N-linked glycoprotein (SIBLING) family proteins. OPN has indeed at least two types of receptors, both broadly expressed in many cell types, but highly upregulated particularly in immune cells. It can interact with integrins via N-terminal domains and CD44 receptors via C-terminal domains. OPN interaction with CD44 causes chemotaxis of neutrophils and macrophages, whereas interactions with integrins relay immune cell spreading and activation signals^65–70^. OPN (SPP1):Integrin pairings were however not detected in our analysis. In the case of the two other predicted significant OPN (SPP1) L-R pairings, SPP1:CCR8 and SPP1:PTGER4 (**Figure 3B and Supplemental Figure 2**) it is unclear whether these interactions produce cellular signals, as these pairings were predicted by CellphoneDB based on protein-protein interaction data curated from various experimental approaches^21,22^, but it is not known whether these interactions produce a cellular signal. Which target cells are impacted by OPN in our model remains to be determined. In our lung scRNAseq dataset CD44 is broadly expressed at low levels in lung stroma, but robustly expressed in immune cells, and highly upregulated by injury in lung neutrophils, monocytes/macrophages, and T cells (data not shown). This suggests that OPN might preferentially target immune cells rather than stromal cells of the lung.

In summary, our work identifies circulating osteopontin released by the injured kidney as a key mediator of lung inflammation and lung edema with respiratory compromise. Our human sample data and available published evidence suggest that therapeutic targeting of OPN or of its associated regulatory or target components should be evaluated in patients with multiorgan failure that are at high risk of acute lung injury or that already have acute lung injury, particularly in the presence of AKI. Any benefit in this area could lead to a meaningful if not substantial reduction of the very high mortality of multiorgan failure.

## Acknowledgements

We are grateful to Sanjay Jain, Michael Rauchman and Peter Herrlich for numerous discussions, insights and reviewing this manuscript. We also thank Kristy Conlon and the Kidney Translational Research Center at Washington University (supported by the Division of Nephrology and P30 DK079333) for providing clinical data and samples from AKI patients. Andreas Herrlich and this work was supported by NIH RO1s DK121200, DK108947 and VA Merit Award BX005322, and Eirini Kefaloyianni was supported by the American Heart Association Career Development Award 20CDA35320006 and the American Society of Nephrology Career Development Award (KidneyCure Carl W. Gottschalk Research Scholar Grant) P21-03591.

## Materials & Methods

### Animals

For all studies, adult (8-12 weeks old) male mice were used in accordance with the animal care and use protocol approved by the Institutional Animal Care and Use Committee of Washington University School of Medicine. C57BL/6J (B6) (JAX stock 000664) and B6.129S6(Cg)-*Spp1^tm1Blh^*/J (*OPN*^-/-^) (JAX stock 004936) were purchased from Jackson Laboratories.

### Surgeries

Bilateral renal ischemia for 20 minutes or 16 minutes (mild AKI) at 37 °C was induced in both kidneys using the flank approach as previously reported by cross-clamping both renal pedicles^71^. Sham operations were performed with exposure of both kidneys, but without induction of ischemia. Syngeneic WT (B6 to B6) or syngeneic KO (*OPN*^-/-^ to B6) kidney transplants were performed as previously described^72^. Anesthesia was induced by a mixture of ketamine (80-100 mg/kg) and xylazine HCl (8-12 mg/kg), intraperitoneally and maintained with 1-2 % isoflurane gas, as required. Briefly, B6 or *OPN^-/-^* donor kidneys were implanted into the abdominal cavity of B6 recipient mice, where the donor suprarenal aorta and renal vein were anastomosed to the recipient infrarenal aorta and inferior vena cava, respectively. The ureter was reconstructed by direct ureter to bladder insertion^73^. Donor kidneys were subjected to 20 min warm ischemia after procurement before they were maintained on ice for implantation, and cold ischemic times were less than 40 minutes. Sham-operated mice underwent the same surgical procedure except for the transplant.

### Single cell preparation for RNA sequencing

Kidneys were minced into small pieces (<1mm^3^), and incubated in tissue dissociation buffer (1 mg/ml Liberase TM, 0.7 mg/ml Hyaluronidase, 80U/ml DNAse in PBS) for 30 min in 37°C. Single cells were released from the digested tissue by pipetting 10 times and the cell suspension was filtered through a 70 µm sieve (Falcon). 10% Fetal Bovine Serum (FBS) was added to stop the enzymatic reaction. Cells were collected by centrifugation (300 g at 4°C for 5 min) and resuspended in red blood cell lysis buffer (155mM NH4Cl, 10mM KHCO3, 1mM EDTA, pH 7.3) for 1 min at room temperature. After washing in PBS, cells were used fresh for analysis by scRNAseq.

### Single Cell RNA Sequencing

Single cell RNA sequencing (scRNAseq) analysis of 4 pooled sham or 4 pooled AKI samples, for both kidney and lung (sham kidney/AKI kidney, sham lung/AKI-ALI lung), was performed as described previously^74^. Briefly, cells were stained with propidium iodide (PI) and live cells were sorted using FACS Aria III (BD Biosciences). Libraries were prepared utilizing the v2 Chromium™ Single Cell 5’ Library Kit and Chromium instrument (10x Genomics, Pleasanton, CA). Full-length cDNA was amplified, and libraries were submitted to Genome Technology Access Center (GTAC) of Washington University in St. Louis for sequencing at a depth of 50,000 reads. All processing steps were performed using Seurat V3. Quality control was first performed on each library to find appropriate filtering thresholds for each. Expression matrices for each sample were loaded into R as Seurat objects, retaining only cells that have more than 200 and less than 3200 genes. Poor quality cells with >10% mitochondrial genes were removed. Any gene not expressed in at least three cells was removed. SCTransform was used for normalization scaling and variance stabilization (https://github.com/ChristophH/sctransform). This was done to reduce bias introduced by technical variation, sequencing depth, and capture efficiency. We assigned an identifier for Sham Kidney, AKI Kidney, Sham Lung, AKI-ALI Lung to tell them apart. Integration of Kidney and Lung single cell data was done using the harmony package (https://github.com/immunogenomics/harmony) to control for batch effects when integrating data from different samples. After QC and integration, 13,882 kidney cells and 15,167 lung cells were further analyzed. We identified 15 clusters in kidney and 20 clusters in lung. We visualized cell clustering using UMAP. To assign cluster identities, we first compiled a list of lung and kidney cell types and their currently established markers^18,19^ and assessed the expression of those markers and additional known canonical markers using the FindAllMarkers () function in Seurat.

### Ligand-Receptor pairing analysis

To infer cell-to-cell communication between kidney and lung cell types from scRNAseq or snRNAseq data we performed Ligand-Receptor pairing analysis using CellPhoneDB^20^ (cellphonedb.org). First, we integrated all data sets from kidney (sham kidney and AKI kidney (4h, 12h, 24)) and lung (sham lung and AKI-ALI lung) using reciprocal PCA as implemented in Seurat. The 4h and 12h AKI kidney snRNAseq data were from published data^18^. CellPhoneDB contains a highly curated set of human protein-protein interactions and protein complexes, thus mouse genes were mapped to their high-confidence human one-to-one ortholog using homology mappings from ENSEMBL. CellPhoneDB statistical analysis was performed with default settings between all kidney and lung cell populations, conditions, and time points simultaneously to increase statistical power. Finally, we considered only co-expressed pairs with a ligand expressed in a kidney cell population and its cognate receptor expressed in a lung cell population with significant cell-type specific co-expression as compared to randomly shuffled cells (p value < 0.01). CellphoneDB calculates an empirical p value of significance, with higher p-values indicating higher significance.

### Histology, Immunofluorescence staining and Quantification

Mice were anesthetized by ketamine cocktail (20 mg/mL ketamine, 2 mg/mL xylazine in 0.9% sodium chloride solution) and then perfused with PBS. Lungs were harvested at the indicated times. Four lobes of the right lung were cut and divided. For histology, 2 lobes (superior and middle lobes) were inserted into 4% paraformaldehyde, located inside a 10 mL syringe, and inflated by creating pressure with the syringe plunger after locking the syringe. After inflation, the lobes were fixed in 4% paraformaldehyde overnight at 4 ℃, and then processed and embedded in paraffin. Sections (4 μm thick) were stained with hematoxylin and eosin. Alveolar wall thickness was measured as previously described using ImageJ^75^. For immunofluorescence, 1 lobe (inferior lobe) of the right lung was placed into PBS, located inside a 10 mL syringe, inflated by creating pressure with the syringe plunger after locking the syringe. Fresh frozen lung sections (7μm) were fixed in 4% paraformaldehyde, permeabilized in 0.1% Triton X-100 for 3 minutes and blocked for 1 hour with 10% normal goat serum supplemented with 1% BSA in PBS. Sections were incubated with primary antibodies in blocking solution overnight at 4 ℃. The primary antibodies used were rat anti-mouse Ly6G/Ly6C (clone RB6-8C5, 14-5931, eBioscience, 1:100), rabbit anti-CD68 (ab125212, abcam, 1:200), rat anti-mouse CD31 (550274, BD Pharmingen^TM^, 1:200), rabbit anti-ZO1 (ab221547, abcam, 1:500), rat anti-Nidogen (sc-33706, Santa Cruz, 1:200). After extensive washes with PBS, fluorescently conjugated secondary antibodies were applied at 1:300 dilution for 1h at room temperature. The secondary antibodies used were as follows: Alexa 594 goat anti-rabbit (A-11037), Alexa 488 goat anti-rat (A-11006). For quantitative analysis, we selected six representative areas which were captured with a Nikon Eclipse E800 microscope at a 200x magnification.

### Electron microscopy

For transmission electron microscopy (TEM), small pieces (1-2 mm cubes) of lung tissue were fixed with 2% paraformaldehyde and 2% glutaraldehyde in 0.1 M sodium cacodylate buffer (pH7.4). Embedding and sectioning were performed by the Washington University Center for Cellular Imaging. To assess the integrity of endothelial cell-cell junction, junction length was measured from digital images using Image J software expressed as a relation of electron dense cortical protein complex area to the total length of cell-cell contact between endothelial cells as previously described^76^.

### Evans blue injection

Mice were injected retro-orbitally with 30 μg/μL Evans blue dye (Sigma-Aldrich) in PBS (50 μg/g of body weight). After 20 minutes, mice were anesthetized and perfused via the left ventricle with 40 mL of PBS, and lungs were removed and additionally rinsed with PBS. The left lung was cut in half and each half was weighed. Evans blue dye was extracted from one half of the lung by incubation with 200 μl formamide (56℃ for 24h) and the concentration of Evans blue was estimated by spectrophotometer (620 nm). The other lung half was dried in an incubator set at 65℃. The dry weight was obtained after 48 h of incubation and the ratio of wet-to-dry weight was calculated. The resulting unit of Evans Blue plasma extravasation was OD_620_/g dry weight^77^.

### OPN-conjugation, injection, and detection

Recombinant Mouse OPN (R&D Systems, 441-OP) was conjugated to Alexa Fluor 555 (Molecular Probes^TM^, A30007) according to the manufacturer. Briefly, OPN protein was reconstituted at 1mg/mL in sterile PBS. One vial of Alexa Fluor 555 succinimidyl ester was dissolved in 10 μl dH_2_O. The reaction mixture containing 25 μl recombinant OPN, 1.66 μl Alexa Fluor 555 and 2.5 μl of 1 M sodium bicarbonate was incubated for 15 minutes at room temperature. After purification, the concentration of the protein was read at 280 and 555 nm by using NanoDrop ND-2000C spectrophotometer (Thermo scientific). Alexa Fluor 555 conjugated OPN was intravenously injected into mice after 20min of bilateral renal ischemia at reperfusion or in sham mice at an equivalent matching time point. Mice were sacrificed 30 minutes or 1 hour later, and mice were anesthetized and perfused via the left ventricle with 40 mL of PBS, before lungs were removed and additionally rinsed with PBS. Lung were then processed to frozen sections as described. and assessed for deposition of OPN into lung tissue by confocal microscopy.

### Human samples

Deidentified human samples of patients with AKI without multiorgan failure, of CKD patients or of healthy controls were provided by the Kidney Translational Research Core at Washington University in St. Louis. Cause of AKI was as follows: 8 acute tubular necrosis (ATN), 1 oxalate crystal ATN, 1 AKI of obstructive etiology, 1 contrast nephropathy, 1 UVJ stone AKI, 1 AKI with HIV and 1 AKI with malignant HTN. Patients did not have clinical evidence of other organ failure based on clinical data review and serum biochemistries.

### Quantitative RT-PCR

Total RNA was isolated from mouse kidneys and lungs using the Direct-zol RNA MiniPrep Plus kit (Cat.No: R2072) following the manufacturers’ instructions. Total RNA was reverse transcribed using the QuantiTect RT kit (Qiagen) and real-time PCR was performed with Fast SYBR Green (Qiagen). Primer sequences are provided in **the table** below. GAPDH was used as the housekeeping gene. Data were analyzed using the delta-delta Ct method.

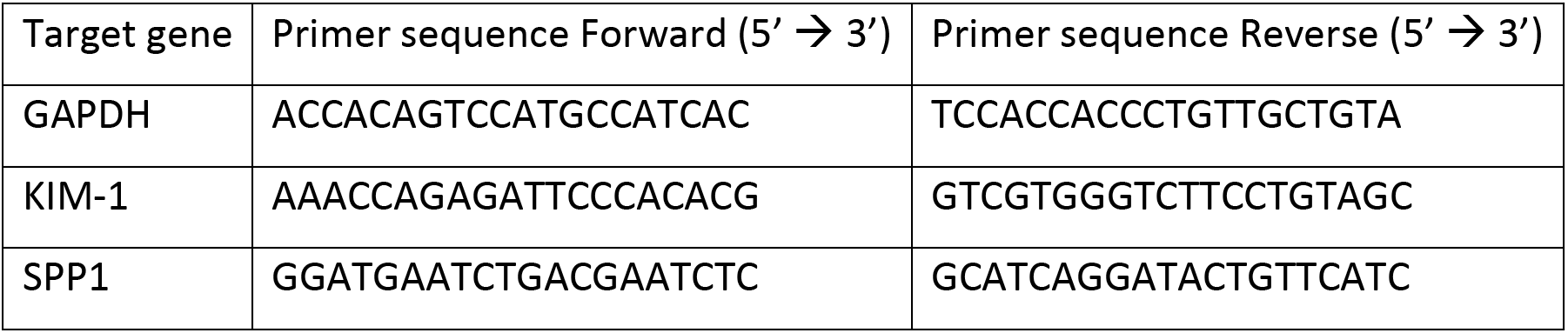

### ELISA

Human OPN, mouse OPN and mouse KIM-1 were measured in human or mouse serum samples using ELISA kits (DY1433, DY441 and DY1817 respectively, all from R&D) as per the manufacturer’s instructions. Serum dilutions for ELISAs were as follows: mouse KIM1-1:10 dilution; mouse OPN: Sham-1:2000, Injured-1:4000 dilution, Sham transplant-1:1000 dilution, WT kidney transplant and OPN-KO kidney transplant-1:2000 dilution; Human serum OPN: Healthy control (1:500), AKI patients (1:1000 dilution), and CKD patients (1:500).

### Mass cytometry CYTOF

Single cell preparations were analyzed by mass cytometry as previously described^78^. Briefly, cells were labeled using a previously validated and titrated antibody cocktail for surface markers (all antibodies conjugated by the manufacturer-Fluidigm) diluted in Fluidigm MaxPar Cell Staining Buffer (CSB) (1 hour at 4 °C). After two washes in CSB, cells were fixed in 2% PFA for 20 min at room temperature, washed, stained with MaxPar Intercalator-IR (Fluidigm), and filtered into cell strainer cap tubes. Data was then acquired on a CyTOF2/Helios instrument (Fluidigm) and analyzed with the CytoBank software using our recently described gating strategy^78^.

### Renal function

Serum creatinine was assessed by an LC-mass spectrometry-based assay at the O’Brien Core Center for Acute Kidney Injury Research (University of Alabama School of Medicine, Birmingham, Alabama, USA). BUN levels were measured using the DiUR100 kit (Thermo Scientific) according to the manufacturer’s instructions.

### Statistical Analysis

Statistical analyses were carried out using Graphpad Prism. Two-tailed, unpaired t tests, one-sample t tests, one-way ANOVA, and two-way ANOVA were used to determine statistical significance in quantification. All results were expressed as mean values ± SD and P values < 0.05 were considered as statistically significant.

## References

1. Zhou X, Franklin RA, Adler M, Jacox JB, Bailis W, Shyer JA, et al.: Circuit Design Features of a Stable Two-Cell System. Cell 172: 744–757.e17, 2018

2. Rouault H, Hakim V: Different Cell Fates from Cell-Cell Interactions: Core Architectures of Two-Cell Bistable Networks. Biophys J 102: 417–426, 2012

3. Bonnans C, Chou J, Werb Z: Remodelling the extracellular matrix in development and disease. Nat Rev Mol Cell Bio 15: 786–801, 2014

4. Bich L, Pradeu T, Moreau J-F: Understanding Multicellularity: The Functional Organization of the Intercellular Space. Front Physiol 10: 1170, 2019

5. Teixeira JP, Ambruso S, Griffin BR, Faubel S: Pulmonary Consequences of Acute Kidney Injury. Seminars in nephrology 39: 3–16, 2019

6. Walcher A, Faubel S, Keniston A, Dennen P: In Critically Ill Patients Requiring CRRT, AKI Is Associated with Increased Respiratory Failure and Death Versus ESRD. Renal Failure 33: 935–942, 2011

7. Faubel S, Edelstein CL: Mechanisms and mediators of lung injury after acute kidney injury. Nature reviews Nephrology 12: 48–60, 2016

8. Bezerra R, Teles F, Mendonca PB, Damte T, Likaka A, Ferrer-Miranda E, et al.: Outcomes of critically ill patients with acute kidney injury in COVID-19 infection: an observational study. Renal Failure 43: 911–918, 2021

9. Darmon M, Clec’h C, Adrie C, Argaud L, Allaouchiche B, Azoulay E, et al.: Acute Respiratory Distress Syndrome and Risk of AKI among Critically Ill Patients. Clin J Am Soc Nephro 9: 1347–1353, 2014

10. Thompson BT, Chambers RC, Liu KD: Acute Respiratory Distress Syndrome. The New England journal of medicine 377: 562–572, 2017

11. Lertjitbanjong P, Thongprayoon C, Cheungpasitporn W, O’Corragain OA, Srivali N, Bathini T, et al.: Acute Kidney Injury after Lung Transplantation: A Systematic Review and Meta-Analysis. J Clin Medicine 8: 1713, 2019

12. Gameiro J, Fonseca JA, Outerelo C, Lopes JA: Acute Kidney Injury: From Diagnosis to Prevention and Treatment Strategies. J Clin Medicine 9: 1704, 2020

13. Lee SA, Cozzi M, Bush EL, Rabb H: Distant Organ Dysfunction in Acute Kidney Injury: A Review. American journal of kidney diseases: the official journal of the National Kidney Foundation 2018

14. Aranda-Valderrama P, Kaynar AM: The Basic Science and Molecular Mechanisms of Lung Injury and Acute Respiratory Distress Syndrome. International anesthesiology clinics 56: 1–25, 2018

15. Doi K, Ishizu T, Tsukamoto-Sumida M, Hiruma T, Yamashita T, Ogasawara E, et al.: The high-mobility group protein B1-Toll-like receptor 4 pathway contributes to the acute lung injury induced by bilateral nephrectomy. Kidney International 86: 316–326, 2014

16. White LE, Cui Y, Shelak CMF, Lie ML, Hassoun HT: Lung endothelial cell apoptosis during ischemic acute kidney injury. Shock (Augusta, Ga.) 38: 320–327, 2012

17. White LE, Santora RJ, Cui Y, Moore FA, Hassoun HT: TNFR1-dependent pulmonary apoptosis during ischemic acute kidney injury. American journal of physiology Lung cellular and molecular physiology 303: L449–59, 2012

18. Kirita Y, Wu H, Uchimura K, Wilson PC, Humphreys BD: Cell profiling of mouse acute kidney injury reveals conserved cellular responses to injury. Proc National Acad Sci 117: 15874–15883, 2020

19. Angelidis I, Simon LM, Fernandez IE, Strunz M, Mayr CH, Greiffo FR, et al.: An atlas of the aging lung mapped by single cell transcriptomics and deep tissue proteomics. Nat Commun 10: 963, 2019

20. Efremova M, Vento-Tormo M, Teichmann SA, Vento-Tormo R: CellPhoneDB: inferring cell–cell communication from combined expression of multi-subunit ligand–receptor complexes. Nat Protoc 15: 1484–1506, 2020

21. Brown KR, Jurisica I: Unequal evolutionary conservation of human protein interactions in interologous networks. Genome Biol 8: R95, 2007

22. Brown KR, Jurisica I: Online Predicted Human Interaction Database. Bioinformatics 21: 2076–2082, 2005

23. Kruger TE, Miller AH, Godwin AK, Wang J: Bone sialoprotein and osteopontin in bone metastasis of osteotropic cancers. Crit Rev Oncol Hemat 89: 330–341, 2014

24. Ramaiah SK, Rittling S: Pathophysiological Role of Osteopontin in Hepatic Inflammation, Toxicity, and Cancer. Toxicol Sci 103: 4–13, 2008

25. Fisher LW, Torchia DA, Fohr B, Young MF, Fedarko NS: Flexible Structures of SIBLING Proteins, Bone Sialoprotein, and Osteopontin. Biochem Bioph Res Co 280: 460–465, 2001

26. Pichler R, Giachelli CM, Lombardi D, Pippin J, Gordon K, Alpers CE, et al.: Tubulointerstitial disease in glomerulonephritis. Potential role of osteopontin (uropontin). Am J Pathology 144: 915–26, 1994

27. Oldberg A, Franzén A, Heinegård D: Cloning and sequence analysis of rat bone sialoprotein (osteopontin) cDNA reveals an Arg-Gly-Asp cell-binding sequence. Proc National Acad Sci 83: 8819–8823, 1986

28. Denhardt DT, Guo X: Osteopontin: a protein with diverse functions. Faseb J 7: 1475–1482, 1993

29. Murugaiyan G, Mittal A, Weiner HL: Increased Osteopontin Expression in Dendritic Cells Amplifies IL-17 Production by CD4+ T Cells in Experimental Autoimmune Encephalomyelitis and in Multiple Sclerosis. J Immunol 181: 7480–7488, 2008

30. Ashkar S, Weber GF, Panoutsakopoulou V, Sanchirico ME, Jansson M, Zawaideh S, et al.: Eta-1 (Osteopontin): An Early Component of Type-1 (Cell-Mediated) Immunity. Science 287: 860–864, 2000

31. Castello LM, Baldrighi M, Molinari L, Salmi L, Cantaluppi V, Vaschetto R, et al.: The Role of Osteopontin as a Diagnostic and Prognostic Biomarker in Sepsis and Septic Shock. Cells 8: 174, 2019

32. Carbone F, Bonaventura A, Vecchiè A, Meessen J, Minetti S, Elia E, et al.: Early osteopontin levels predict mortality in patients with septic shock. Eur J Intern Med 78: 113–120, 2020

33. Thompson BT, Chambers RC, Liu KD: Acute Respiratory Distress Syndrome. The New England journal of medicine 377: 1904–1905, 2017

34. Vassiliou AG, Kotanidou A, Dimopoulou I, Orfanos SE: Endothelial Damage in Acute Respiratory Distress Syndrome. Int J Mol Sci 21: 8793, 2020

35. Dolmatova EV, Wang K, Mandavilli R, Griendling KK: The effects of sepsis on endothelium and clinical implications. Cardiovasc Res 2020

36. Hotchkiss RS, Moldawer LL, Opal SM, Reinhart K, Turnbull IR, Vincent J-L: Sepsis and septic shock. Nat Rev Dis Primers 2: 16045, 2016

37. Prescott HC, Angus DC: Enhancing Recovery From Sepsis: A Review. Jama 319: 62–75, 2018

38. Michalick L, Weidenfeld S, Grimmer B, Fatykhova D, Solymosi PD, Behrens F, et al.: Plasma mediators in patients with severe COVID-19 cause lung endothelial barrier failure. Eur Respir J 57: 2002384, 2021

39. Klein CL, Hoke TS, Fang W-F, Altmann CJ, Douglas IS, Faubel S: Interleukin-6 mediates lung injury following ischemic acute kidney injury or bilateral nephrectomy. Kidney International 74: 901–909, 2008

40. Kim DJ, Park SH, Sheen MR, Jeon US, Kim SW, Koh ES, et al.: Comparison of experimental lung injury from acute renal failure with injury due to sepsis. Respiration; international review of thoracic diseases 73: 815–824, 2006

41. Kramer AA, Postler G, Salhab KF, Mendez C, Carey LC, Rabb H: Renal ischemia/reperfusion leads to macrophage-mediated increase in pulmonary vascular permeability. Kidney International 55: 2362–2367, 1999

42. Ministrini S, Carbone F, Montecucco F: Emerging role for the inflammatory biomarker osteopontin in adverse cardiac remodeling. Biomark Med 14: 1303–1306, 2020

43. Martinez SR, Fernández DI, Cuesta VMM, Genre F, Pulito-Cueto V, Rozas SMF, et al.: Osteopontin and interleukin-6 as biomarkers of interstitial lung disease. 2717, 2020

44. Varım C, Demirci T, Cengiz H, Hacıbekiroğlu İ, Tuncer FB, Çokluk E, et al.: Relationship between serum osteopontin levels and the severity of COVID-19 infection. Wien Klin Wochenschr 133: 298–302, 2021

45. Loosen SH, Roderburg C, Kauertz KL, Pombeiro I, Leyh C, Benz F, et al.: Elevated levels of circulating osteopontin are associated with a poor survival after resection of cholangiocarcinoma. J Hepatol 67: 749–757, 2017

46. Behnes M, Brueckmann M, Lang S, Espeter F, Weiss C, Neumaier M, et al.: Diagnostic and prognostic value of osteopontin in patients with acute congestive heart failure. Eur J Heart Fail 15: 1390–1400, 2013

47. Lorenzen JM, Hafer C, Faulhaber-Walter R, Kümpers P, Kielstein JT, Haller H, et al.: Osteopontin predicts survival in critically ill patients with acute kidney injury. Nephrology, dialysis, transplantation : official publication of the European Dialysis and Transplant Association - European Renal Association 26: 531–7, 2010

48. Rosenberg M, Zugck C, Nelles M, Juenger C, Frank D, Remppis A, et al.: Osteopontin, a New Prognostic Biomarker in Patients With Chronic Heart Failure. Circulation Hear Fail 1: 43–49, 2008

49. Cappellano G, Abreu H, Raineri D, Scotti L, Castello L, Vaschetto R, et al.: High levels of circulating osteopontin in inflammatory lung disease regardless of Sars-CoV-2 infection. Embo Mol Med e14124, 2021

50. Gibellini L, Biasi SD, Paolini A, Borella R, Boraldi F, Mattioli M, et al.: Altered bioenergetics and mitochondrial dysfunction of monocytes in patients with COVID-19 pneumonia. Embo Mol Med 12: e13001, 2020

51. Roderburg C, Benz F, Cardenas DV, Lutz M, Hippe H-J, Luedde T, et al.: Persistently elevated osteopontin serum levels predict mortality in critically ill patients. Critical care (London, England) 19: 271, 2015

52. Vaschetto R, Nicola S, Olivieri C, Boggio E, Piccolella F, Mesturini R, et al.: Serum levels of osteopontin are increased in SIRS and sepsis. Intens Care Med 34: 2176–2184, 2008

53. Owens E, Tan K-S, Ellis R, Vecchio SD, Humphries T, Lennan E, et al.: Development of a Biomarker Panel to Distinguish Risk of Progressive Chronic Kidney Disease. Biomed 8: 606, 2020

54. Jin Z-K, Tian P-X, Wang X-Z, Xue W-J, Ding X-M, Zheng J, et al.: Kidney injury molecule-1 and osteopontin: New markers for prediction of early kidney transplant rejection. Mol Immunol 54: 457–464, 2013

55. Hirano Y, Aziz M, Yang W-L, Ochani M, Wang P: Neutralization of Osteopontin Ameliorates Acute Lung Injury Induced by Intestinal Ischemia-Reperfusion. Shock 46: 431–438, 2016

56. Zhang J, Yang G, Zhu Y, Peng X, Li T, Liu L: Relationship of Cx43 regulation of vascular permeability to osteopontin-tight junction protein pathway after sepsis in rats. American journal of physiology. Regulatory, integrative and comparative physiology 314: R1–R11, 2017

57. Zhang J, Yang G, Zhu Y, Peng X, Li T, Liu L: Role of connexin 43 in vascular hyperpermeability and relationship to Rock1-MLC20 pathway in septic rats. Am J Physiol-lung C 309: L1323–L1332, 2015

58. Moschos C, Porfiridis I, Psallidas I, Kollintza A, Stathopoulos GT, Papiris SA, et al.: Osteopontin is upregulated in malignant and inflammatory pleural effusions. Respirology 14: 716–722, 2009

59. Psallidas I, Stathopoulos GT, Maniatis NA, Magkouta S, Moschos C, Karabela SP, et al.: Secreted phosphoprotein-1 directly provokes vascular leakage to foster malignant pleural effusion. Oncogene 32: 528–535, 2013

60. Black RA, Rauch CT, Kozlosky CJ, Peschon JJ, Slack JL, Wolfson MF, et al.: A metalloproteinase disintegrin that releases tumour-necrosis factor-alpha from cells. Nature [Internet] 385: 729–733, 1997 Available from: http://www.nature.com/nature/journal/v385/n6618/abs/385729a0.html

61. Hernando AA, Okamura K, Bhargava R, Kiekhaefer CM, Soranno D, Kirkbride-Romeo LA, et al.: Circulating IL-6 upregulates IL-10 production in splenic CD4+ T cells and limits acute kidney injury-induced lung inflammation. Kidney International 91: 1057–1069, 2017

62. Hassoun HT, Grigoryev DN, Lie ML, Liu M, Cheadle C, Tuder RM, et al.: Ischemic acute kidney injury induces a distant organ functional and genomic response distinguishable from bilateral nephrectomy. American journal of physiology Renal physiology 293: F30–40, 2007

63. Hoke TS, Douglas IS, Klein CL, He Z, Fang W, Thurman JM, et al.: Acute renal failure after bilateral nephrectomy is associated with cytokine-mediated pulmonary injury. Journal of the American Society of Nephrology : JASN 18: 155–164, 2007

64. Hernando AA, Dursun B, Altmann C, Ahuja N, He Z, Bhargava R, et al.: Cytokine production increases and cytokine clearance decreases in mice with bilateral nephrectomy. Nephrology Dialysis Transplantation 27: 4339–4347, 2012

65. Wei J, Marisetty A, Schrand B, Gabrusiewicz K, Hashimoto Y, Ott M, et al.: Osteopontin mediates glioblastoma-associated macrophage infiltration and is a therapeutic target. J Clin Invest 129: 137–149, 2018

66. Zhu B, Suzuki K, Goldberg HA, Rittling SR, Denhardt DT, McCulloch CAG, et al.: Osteopontin modulates CD44-dependent chemotaxis of peritoneal macrophages through G-protein-coupled receptors: Evidence of a role for an intracellular form of osteopontin. J Cell Physiol 198: 155–167, 2004

67. Koh A, Silva APBD, Bansal AK, Bansal M, Sun C, Lee H, et al.: Role of osteopontin in neutrophil function. Immunology 122: 466–475, 2007

68. Hirano Y, Aziz M, Yang W-L, Wang Z, Zhou M, Ochani M, et al.: Neutralization of osteopontin attenuates neutrophil migration in sepsis-induced acute lung injury. Crit Care 19: 53, 2015

69. Zhou Y, Yao Y, Shen L, Zhang J, Zhang JH, Shao A: Osteopontin as a candidate of therapeutic application for the acute brain injury. J Cell Mol Med 24: 8918–8929, 2020

70. Moorman HR, Poschel D, Klement JD, Lu C, Redd PS, Liu K: Osteopontin: A Key Regulator of Tumor Progression and Immunomodulation. Cancers 12: 3379, 2020

71. Yang L, Besschetnova TY, Brooks CR, Shah JV, Bonventre JV: Epithelial cell cycle arrest in G2/M mediates kidney fibrosis after injury. Nature Medicine 16: 535–43, 1p following 143, 2010

72. Kalina SL, Mottram PL: A MICROSURGICAL TECHNIQUE FOR RENAL TRANSPLANTATION IN MICE. Aust Nz J Surg 63: 213–216, 1993

73. Han W, Murray-Segal LJ, Mottram PL: Modified technique for kidney transplantation in mice. Microsurg 19: 272–274, 1999

74. Williams JW, Winkels H, Durant CP, Zaitsev K, Ghosheh Y, Ley K: Single Cell RNA Sequencing in Atherosclerosis Research. Circ Res 126: 1112–1126, 2020

75. Pua ZJ, Stonestreet BS, Cullen A, Shahsafaei A, Sadowska GB, Sunday ME: Histochemical Analyses of Altered Fetal Lung Development Following Single vs Multiple Courses of Antenatal Steroids. J Histochem Cytochem 53: 1469–1479, 2005

76. Hakanpaa L, Kiss EA, Jacquemet G, Miinalainen I, Lerche M, Guzmán C, et al.: Targeting β1-integrin inhibits vascular leakage in endotoxemia. Proc National Acad Sci 115: 201722317, 2018

77. Wick MJ, Harral JW, Loomis ZL, Dempsey EC: An Optimized Evans Blue Protocol to Assess Vascular Leak in the Mouse. J Vis Exp 2018

78. Brody SL, Gunsten SP, Luehmann HP, Sultan DH, Hoelscher M, Heo GS, et al.: Chemokine Receptor 2–targeted Molecular Imaging in Pulmonary Fibrosis. A Clinical Trial. Am J Resp Crit Care 203: 78–89, 2021

