## Supplemental Figures for "Circulating osteopontin released by injured kidneys causes pulmonary inflammation and edema"

### Supplemental Figure 1

**A** L-R pairing Kidney Immune to Lung Immune cells

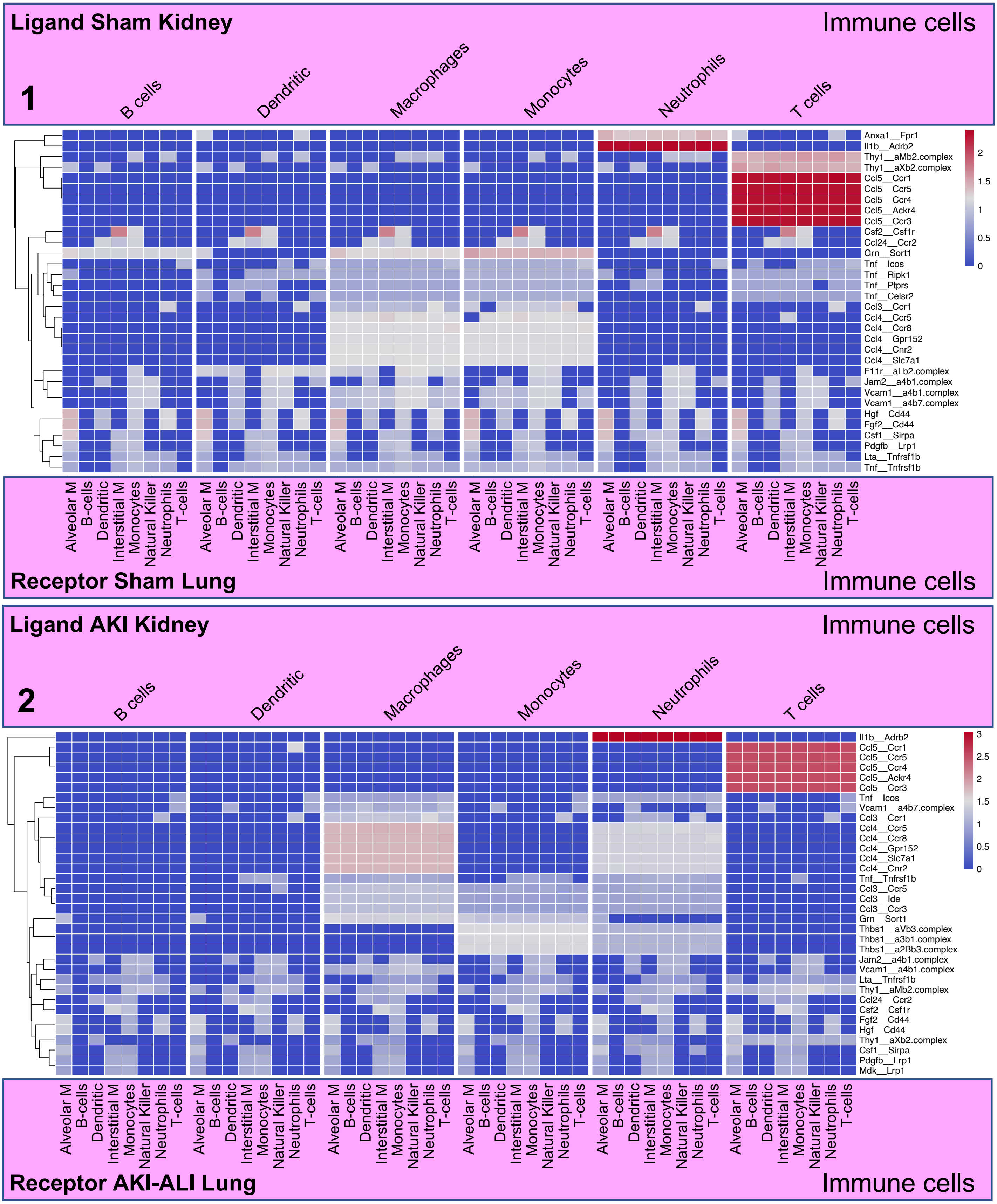

### Supplemental Figure 1 cont.

**B** L-R pairing Kidney Immune to Lung Non-immune cells

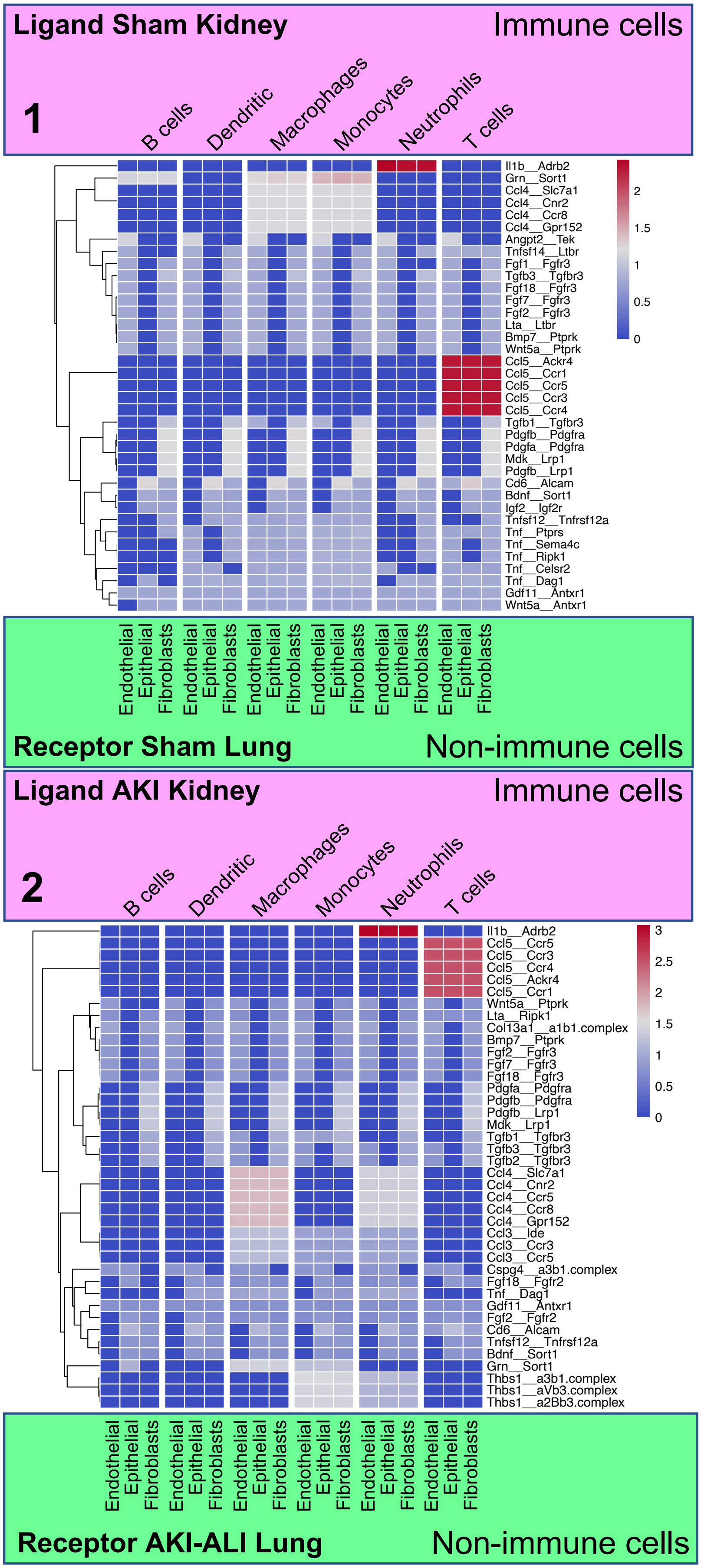

**Supplemental Figure 2**

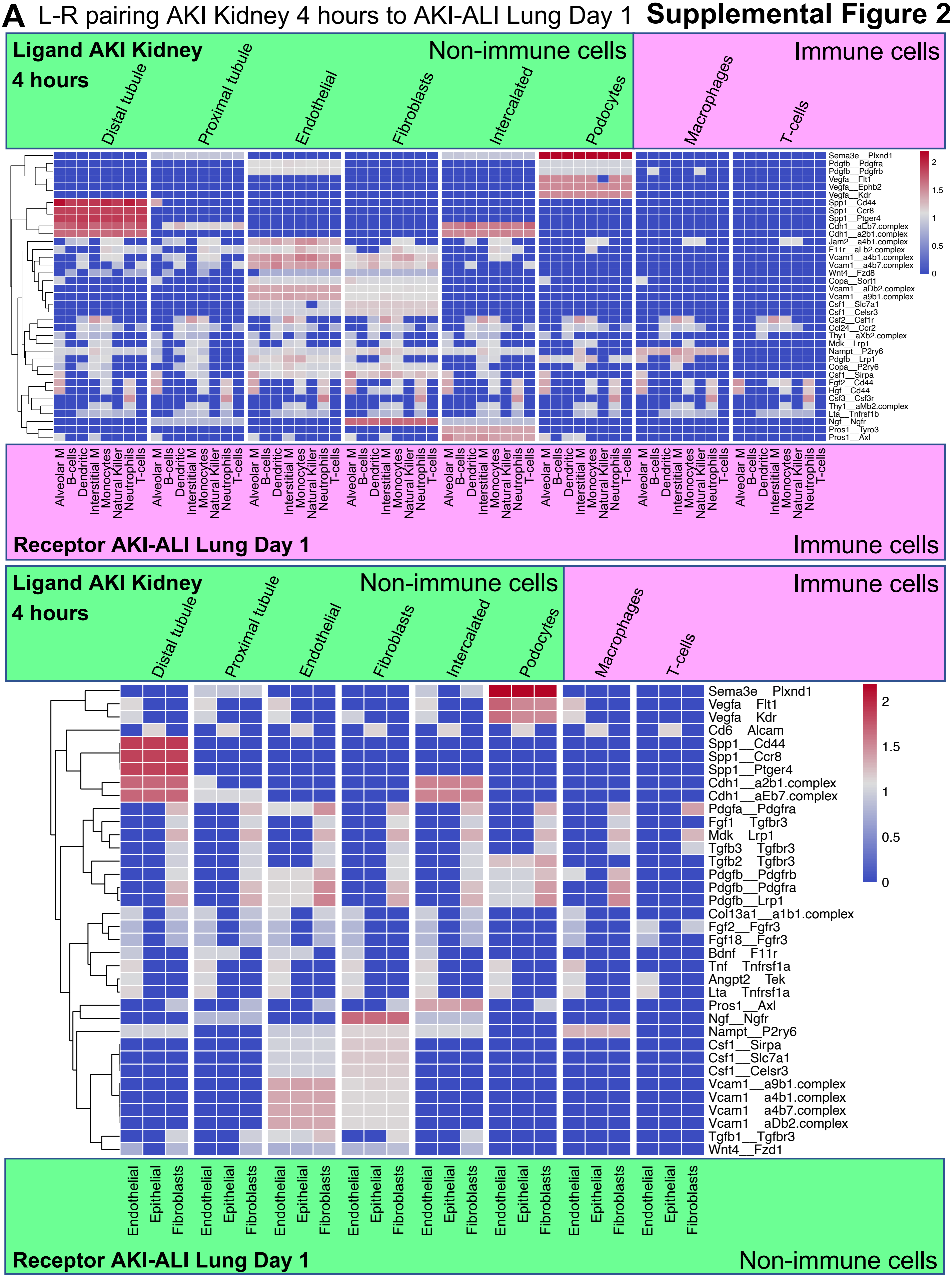

**Supplemental Figure 2**

**B** L-R pairing AKI Kidney 12 hours to AKI-ALI Lung Day 1

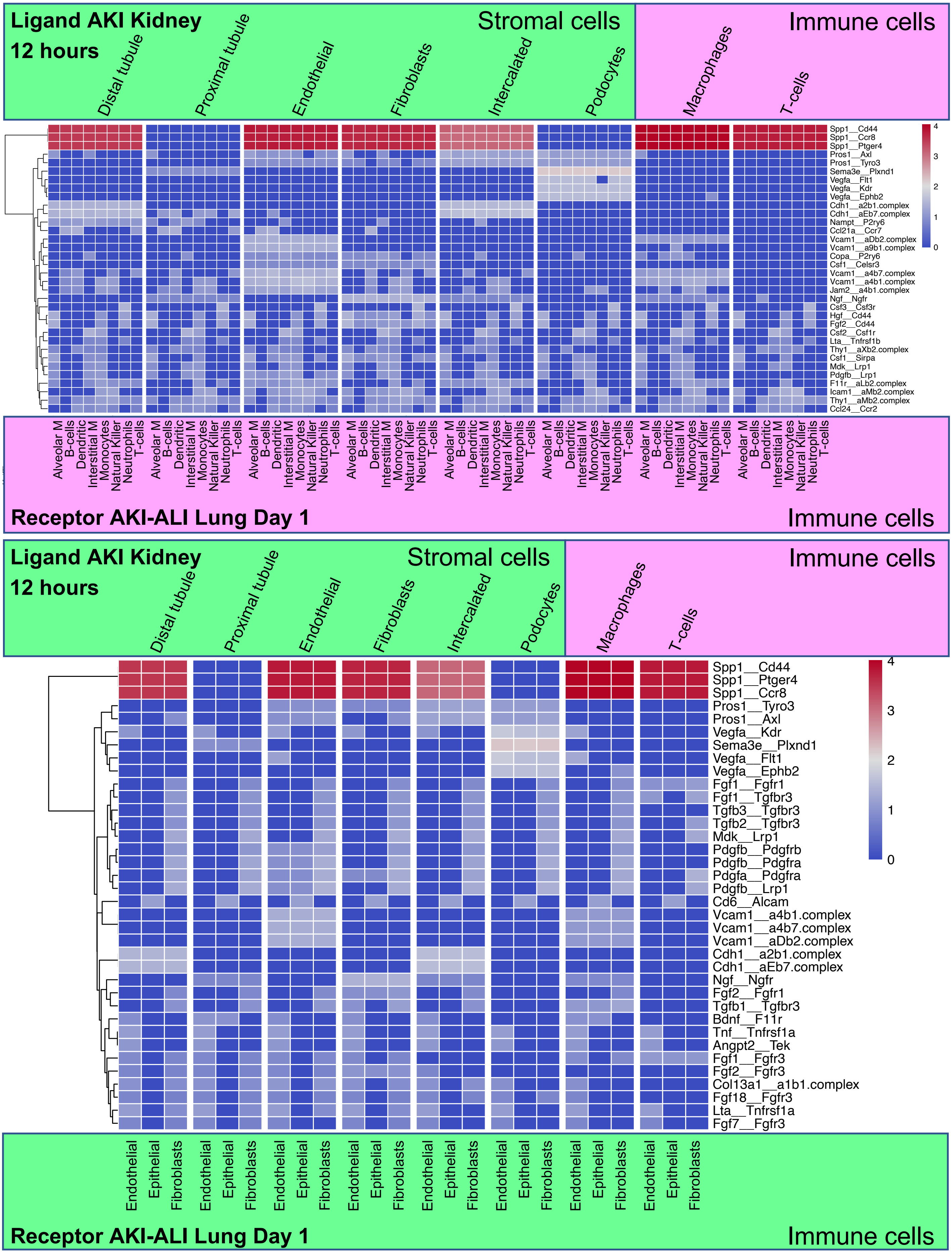

### Supplemental Figure 3

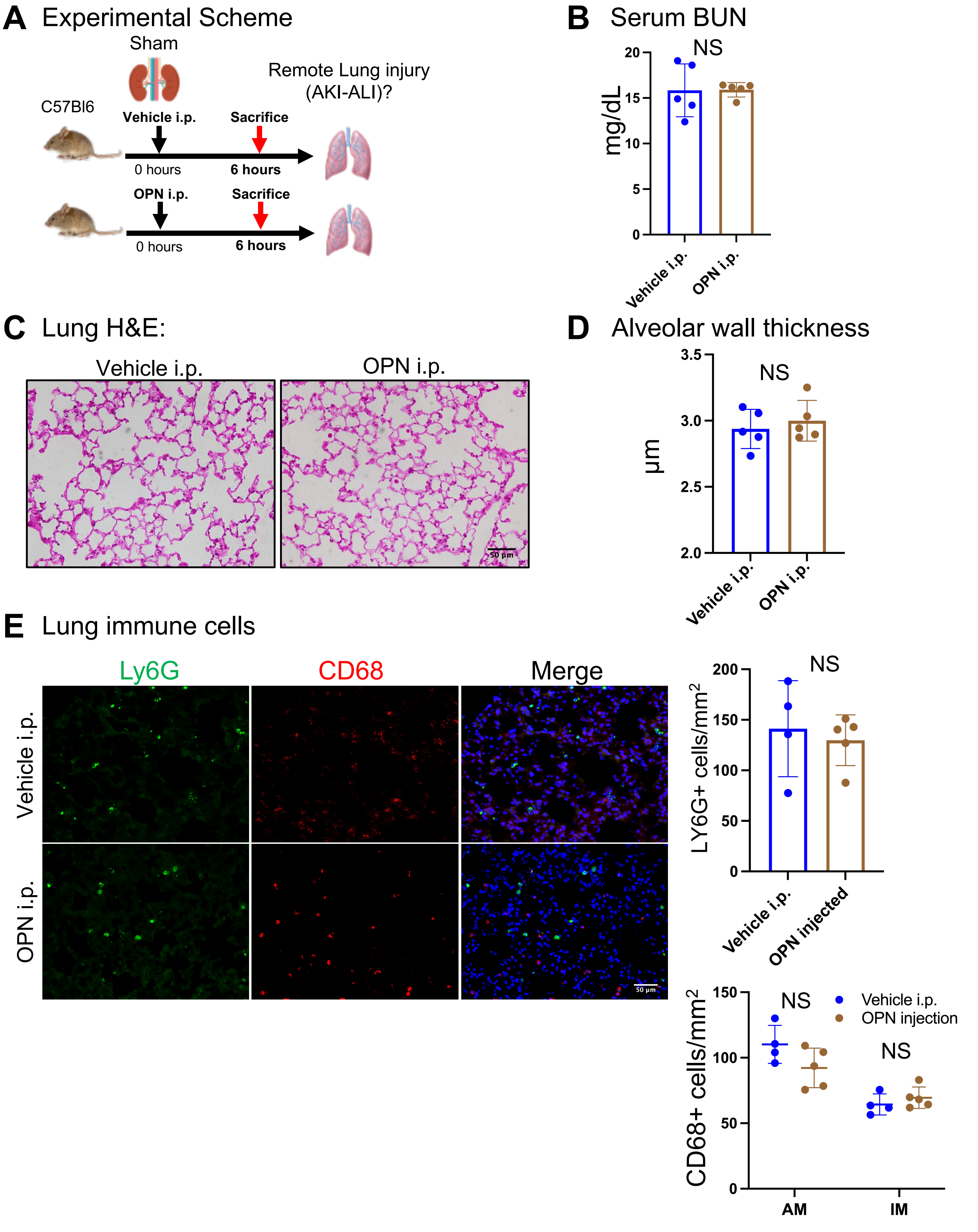
