## Supplemental Figure Legends for "Circulating osteopontin released by injured kidneys causes pulmonary inflammation and edema"

**Supplemental Figure 1: Ligand-Receptor pairing analysis for immune cells** (**A.**) L-R pairing analysis kidney immune cells to lung immune cells for Day 1 after sham (panel 1) or AKI (panel 2), (**B.**) L-R pairing analysis kidney immune cells to lung non-immune cells for Day 1 after sham (panel 1) or AKI (panel 2). p values shown are empirical p values calculated by CellphoneDB, higher values indicate higher L-R pairing significance.

**Supplemental Figure 2: L-R pairing analysis: Integration of published snRNAseq data of the kidney 4hours and 12 hours after AKI with our Day 1 lung scRNAseq data** (**A.**) L-R pairing analysis kidney non-immune or immune cells 4 hours after AKI to lung non-immune or immune cells Day 1 after AKI, (**B.**) L-R pairing analysis kidney non-immune or immune cells 12 hours after AKI to lung non-immune or immune cells Day 1 after AKI. p values shown are empirical p values calculated by CellphoneDB, higher values indicate higher L-R pairing significance. snRNAseq data were from Kirita et al. Proceedings of the National Academy of Sciences 2020.

**Supplemental Figure 3: OPN injection into wt mice without kidney injury does not cause AKI-ALI.** (**A.**) Experimental scheme: *wt* mice are evaluated 6 hours after injection with vehicle control or OPN protein (no AKI injury), (**B.**) Serum BUN 6 hours after vehicle or OPN injection, (**C.**) Lung H&E stain 6 hours after vehicle or OPN injection, (**D.**) Alveolar thickness measurements 6 hours after vehicle or OPN injection, (**E.**) Lung neutrophils (Ly6G$^{+}$, green), alveolar (CD68$^{high}$, large, red) and interstitial macrophages (CD68$^{low}$, small, red) and quantification 6 hours after vehicle or OPN injection, DAPI stain (blue) was used to visualize nuclei. n=4-5 animals per measurement. NS=not significant.
